# Control of spatiotemporal activation of organ-specific fibers in the vagus nerve by intermittent interferential current stimulation

**DOI:** 10.1101/2024.10.22.619669

**Authors:** Nicolò Rossetti, Weiguo Song, Philipp Schnepel, Naveen Jayaprakash, Dimitrios A. Koutsouras, Mark Fichman, Jason Wong, Todd Levy, Mohamed Elgohary, Khaled Qanud, Alice Giannotti, Mary F Barbe, Frank Liu Chen, Geert Langereis, Timir Datta-Chaudhuri, Vojkan Mihajlović, Stavros Zanos

## Abstract

Vagus nerve stimulation (VNS) is emerging as potential treatment for several chronic diseases, however, limited control of fiber activation to promote desired effects over side effects restricts clinical translation. Here we describe a new VNS method that relies on intermittent, interferential sinusoidal current stimulation (i^2^CS) through implanted, multi-contact epineural cuffs. In swine, i^2^CS elicits specific nerve potentials and end organ responses, distinct from equivalent non-interferential sinusoidal stimulation. Comparing experimental results with anatomical trajectories of nerve fascicles from end organs to the stimulation electrode indicates that i^2^CS activates organ-specific fascicles rather than the entire nerve. Experimental results and anatomically realistic, physiologically validated biophysical models of the vagus nerve demonstrate that i^2^CS reduces fiber activation at the focus of interference. Current steering and repetition frequency determine spatiotemporal pattern of vagal fiber activation, allowing tunable and precise control of neural and organ responses. In experiments in a cohort of anesthetized swine, i^2^CS has improved selectivity for a desired effect, mediated by smaller bronchopulmonary fibers, over a side effect, mediated by larger laryngeal fibers, compared to non-interferential sinusoidal or square pulse VNS.

## Introduction

Neural signaling through the vagus nerve is essential for physiological homeostasis through autonomic reflexes (Jänig, 2022) and is implicated in the pathogenesis of several chronic brain, cardiopulmonary, gastrointestinal and inflammatory diseases (Karemaker, 2022). For those reasons, the vagus nerve has emerged as a target for therapeutic neuromodulation, with vagus nerve stimulation (VNS) therapies approved for epilepsy and depression (Afra et al., 2021; Ben-Menachem, 2001; Sackeim et al., 2001) and currently tested in stroke rehabilitation (Dawson et al., 2021), Alzheimer’s disease (Merrill et al., 2006), pain (Chakravarthy et al., 2015), anxiety (George et al., 2008), tinnitus (Tyler et al., 2017), rheumatoid arthritis (Koopman et al., 2016), inflammatory bowel disease (Sinniger et al., 2020), heart failure (De Ferrari et al., 2017), diabetes (Meyers et al., 2016), obesity (Pardo et al., 2007) and pulmonary hypertension (Ntiloudi et al., 2019; Zafeiropoulos et al., 2024a).

Vagus nerve stimulation (VNS) is typically delivered through epineural cuff electrodes implanted around the cervical nerve trunk (Figure 1, B), where sensory and motor fibers travel inside nerve fascicles (Jayaprakash et al., 2023). Small size vagal fibers, such as cholinergic and adrenergic fibers innervating the heart, lungs or abdominal viscera, spatially organized in specific fascicles of the cervical vagus nerve (Jayaprakash et al., 2023) are often responsible for clinically beneficial responses and the actual therapeutic targets of VNS. Currently used VNS devices activate mostly larger fibers, innervating organs like the larynx and pharynx, resulting in side effects which may lead to reduced therapeutic efficacy (Zafeiropoulos et al., 2024b). More spatially-selective VNS delivered through multi-contact cuff electrodes (MCEs), activates organ-specific fibers asymmetrically and differentially (Figure 1, C) (Jayaprakash et al., 2023; Thompson et al., 2024). However, even with MCEs, functional selectivity is limited: larger fibers are still activated before smaller fibers, and fibers located at the periphery of the trunk, closer to the stimulation contacts, are activated before those at the interior of the trunk. Despite its potential translational significance, spatial and temporal control of the activation of vagal fibers, for example of fibers mediating specific desired effects vs. side effects, is currently non feasible, even with the latest MCEs and state-of-the-art stimulation paradigms.

**Figure 1:**
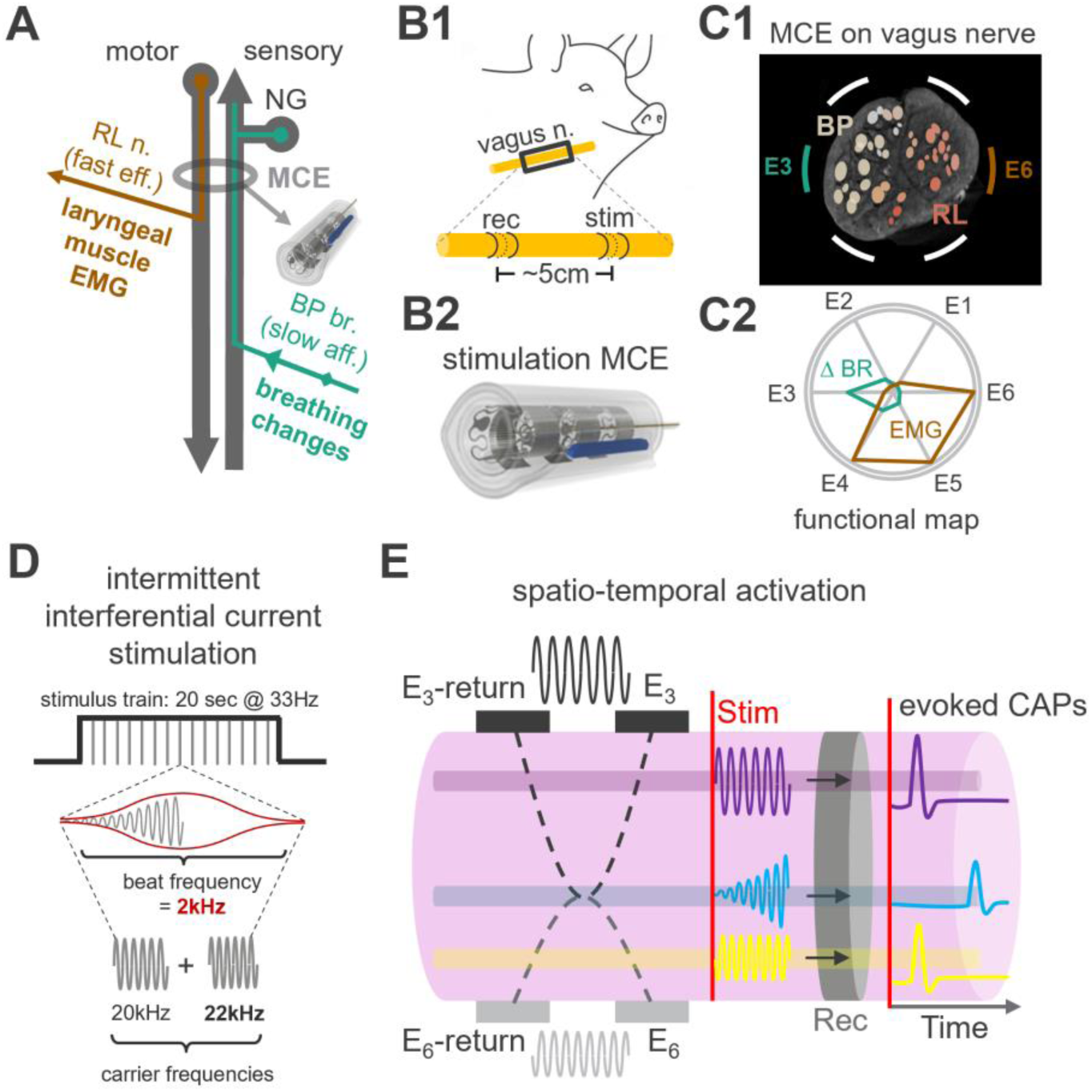
Anatomical basis for control of spatiotemporal activation of fibers by vagus nerve stimulation (VNS) using intermittent, interferential current stimulation (i^2^CS). (A) Schematic of the vagal trunk at the cervical level, below the nodose ganglion (NG), where a multi-contact cuff electrode (MCE) is implanted; shown are fast, motor fibers projecting to laryngeal muscles through the recurrent laryngeal (RL) branch, whose activation produces a laryngeal electromyography (EMG) signal; also, slower, sensory fibers from the lungs, entering the trunk through the bronchopulmonary (BP) branch, whose activation slows down breathing. (B1) Stimulation and recording electrodes placed on the cervical VN of swine used to record evoked nerve potentials and directly assess fiber activation. (B2) Layout of the cylindrical MCE used for VNS, comprising 3 rows of contacts, with 6 contacts in each row. (C1) Schematic cross section of a swine cervical VN with fascicles; fascicle color represents the varying percentages of RL (red) and BP fibers (yellow), determined via post-mortem imaging and fascicle tracking. (C2) Functional map of nerve trunk inferred by single-contact stimulation and recording of physiological responses; contact E3, which is close to BP fascicles, is associated with a strong breathing response (green trace), whereas contact E6, which is close to RL fascicles, is associated with a strong laryngeal EMG response (red trace). Stimulation from other contacts elicits physiological responses with varying magnitudes. (D) I^2^CS waveform in a 20 sec-long stimulus train, with pulse repetition frequency of 33 Hz. Each “pulse” is generated by sinusoidal stimuli with slightly different carrier frequencies (20 and 22 kHz), delivered through separate contacts, which result in amplitude modulation of the short bursts with a beat frequency of 2 kHz (red) through temporal interference. (E) Illustration of the delivery of 2 high frequency sinusoidal stimuli, one between contact E3 and E3-return, and one between contact E6 and E6-return, to produce interference at a specific location inside the nerve trunk. Points close to contacts E3 and E6 do not experience interference or electric field amplitude modulation (AM); the respective fibers (purple and yellow) are activated immediately upon onset of stimulation, resulting in relatively large evoked compound action potentials (CAPs) with short latencies. Points at the focus of interference experience field amplitude modulation, and the respective fibers (blue) are activated to a lesser degree and only after a delay, resulting in smaller evoked CAPs with longer latencies.

Interferential current stimulation (iCS) has recently (re)gained attention as a method for targeted neuromodulation (Grossman et al., 2017; Mirzakhalili et al., 2020). Applying iCS on the brain assumes independent electrical sources with slightly different high frequencies (in the order of kHz) placed outside of the brain result in spatially focused activation of neurons located deeper in the brain, by means of temporal interference and low frequency (tens of Hz) amplitude modulation (Acerbo et al., 2022; von Conta et al., 2021). Whether iCS has a role in selective stimulation of fascicular nerves in general and of the vagus nerve in particular is not known (Botzanowski et al., 2022; Budde et al., 2023). In this paper, we report a novel method for VNS, termed intermittent interferential current stimulation (i^2^CS), that attains spatial and temporal control of activation of organ-specific fibers inside the vagal trunk. Using in vivo experiments in swine and in silico computational modeling, we demonstrate that i^2^CS activates organ-specific fibers in a predictable, spatially focused and temporally precise manner and has improved selectivity for a desired effect over a side effect compared to equivalent, non-interfering sinusoidal and to standard, square pulse VNS.

## Results

### 1. Bronchopulmonary- and laryngeal-specific vagal fibers progressively mix inside nerve fascicles, resulting in a bimodal anatomical organization at the cervical vagus nerve

Use of spatially-selective VNS to preferentially activate a desired effect, e.g., from the lungs, over a side effect, e.g., from laryngeal muscles, relies on anatomical separation between bronchopulmonary- and laryngeal-specific vagal fibers at the site of electrode implantation. Separation of organ-specific fibers inside fascicles at the cervical vagus nerve has been qualitatively demonstrated (Jayaprakash et al., 2023) but has not been quantified, and therefore the anatomical constraints on spatially-selective VNS are unknown. To quantify the anatomical separation of fibers, we tracked the longitudinal trajectories of bronchopulmonary (BP) and recurrent laryngeal (RL) fascicles from the level of branch emergence to the cervical region, identified merges and splits of fascicles, and estimated the mixing of fibers inside fascicles at different levels (Figure 2). We found that at branch emergence and for a few centimeters in the rostral direction, BP- and RL-specific fascicles are spatially almost completely separated from other fascicles (Figure 2, B and 2, C, respectively). However, at more rostral levels, BP, RL and other fascicles progressively merge, and, at the cervical region, no fascicles have fibers originating solely from a single organ; instead fascicles contain a mix of BP, RL and other fibers (Figure 2, D). Despite significant fiber mixing, BP and RL fibers show a distinct spatial arrangement, with BP-rich fascicles in one area of the nerve and RL-rich fascicles in another area (Figure 2, D3, D4), resulting in a bimodal distribution of fibers along a transverse axis (Figure 2, D5).

**Figure 2:**
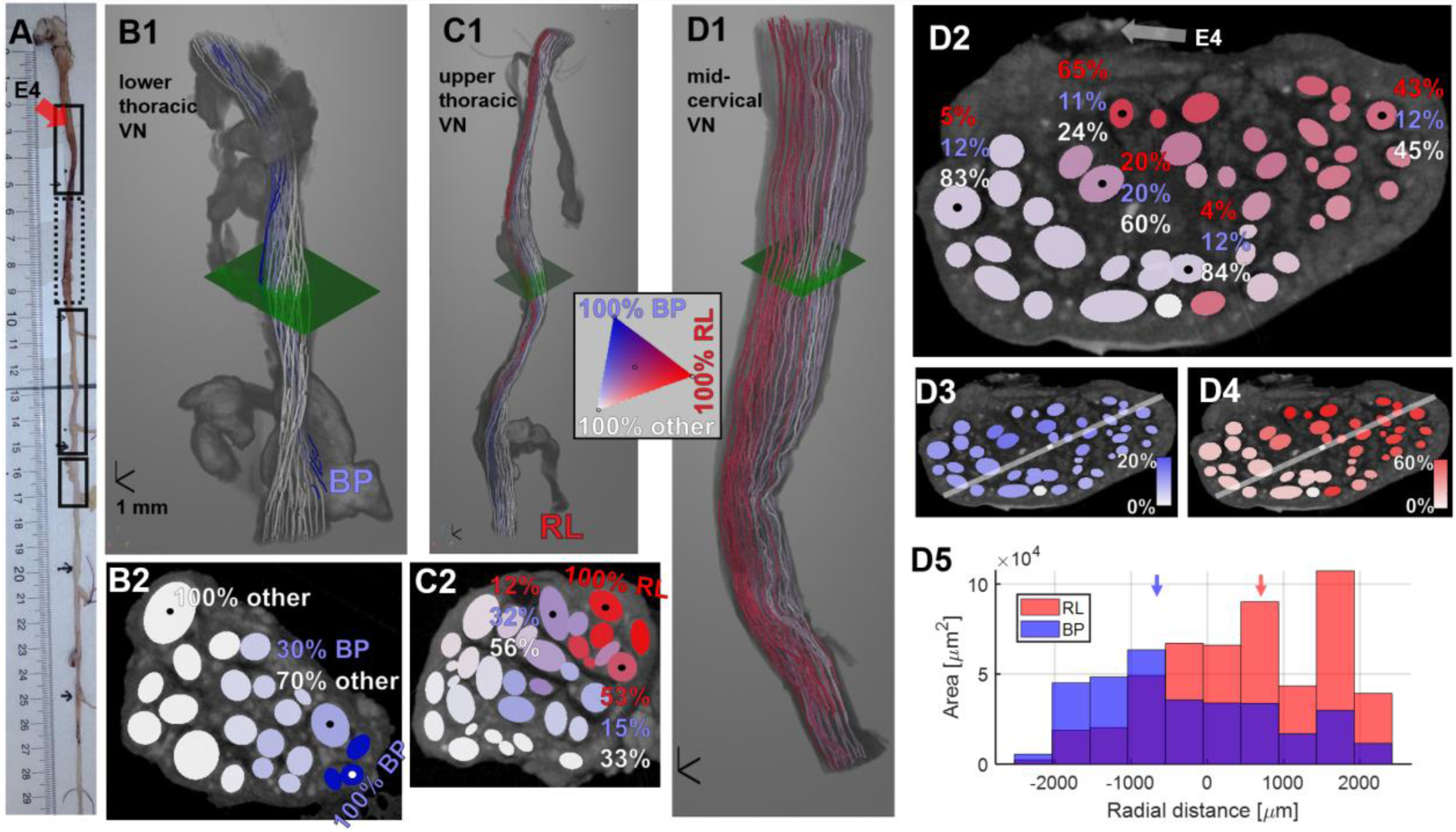
Bronchopulmonary (BP)- and recurrent laryngeal (RL)-specific fibers progressively mix inside nerve fascicles and give rise to a bimodal anatomical organization at the cervical level. (A) After completion of in vivo experiments, the stimulated nerve, along with the RL and BP branches, was dissected, between the nodose ganglion (rostral) and the lower thoracic region (caudal); the exact location of one of the MCE contacts (E4) was marked on the epineurium of the mid-cervical VN with a suture. Each of several segments of the vagal trunk (black rectangles) was imaged with microcomputed tomography (micro-CT), as described previously (Jayaprakash et al., 2023). In the micro-CT data, organ-specific fascicles were tracked longitudinally from branch emergence to the mid-cervical level, fascicle splits and merges were identified and percentages of organ-specific fibers in the resulting fascicle(s) were updated according to relative cross-sectional areas of parent and daughter fascicles. (B1) Reconstructed lower thoracic segment with BP branch emergence and respective fascicles shown in blue. (B2) Cross-section of the vagal trunk shown in B1 (green plane); each fascicle colored according to the percentage of BP fibers. “Other” vagal fibers are those innervating the heart, esophagus and abdominal organs. (C1) Same as B1, for an upper thoracic segment, at RL branch emergence, with respective fascicles shown in red. (C2) Fascicular map at the level of the green plane in C1. Fascicles contain varying numbers of BP, RL and other fibers, represented using a 3-color scale (inset). (D1) Mid-cervical segment, where MCE was implanted. (D2) Fascicular map at level of green plane in D1, with location of MCE contact E4 indicated by the suture marking. (D3) Same map as D2, with colormap corresponding to the % of BP fibers inside fascicles, normalized between maximum and minimum. Diagonal line approximately corresponds to the radial direction defined by 2 of the contacts of the MCE used for i^2^CS in preceding in vivo experiments (E2 and E6). (D4) Percentage of RL fibers (normalized). (D5) Distribution of estimated BP and RL fiber counts projected on the E2-E6 diagonal line, at different distances from the center of the line; blue and red vertical arrows represent the median values of the BP and RL distance distributions (−593 and 547 μm, respectively; p<1^−10^, Wilcoxon rank-sum test).

*Using quantitative anatomical tracking, we document bimodal anatomical organization of organ-specific fibers in the cervical vagus nerve, suggesting that focal stimulation along a transverse direction could improve selectivity of VNS*.

### 2. Interferential stimulation elicits distinct experimental nerve and organ responses that are different than those of sinusoidal stimulation

Interfering current sources give rise to electric fields and amplitude modulations (AM) at distinct spatial locations that are different than those with equivalent non-interfering stimulation (Supplementary Figure S 1) (Mirzakhalili et al., 2020). To test whether interferential VNS activates different areas inside the vagal trunk, thereby engaging different fiber populations, i^2^CS was delivered through pairs of contacts of an MCE; then, evoked compound action potentials (eCAPs) and physiological responses from the lungs and laryngeal muscles were measured. i^2^CS with uneven stimulus intensities results in AM on one side of the nerve (negative steering ratio; Figure 3, A). The fast-fiber eCAP and the respective, fast-fiber-mediated, laryngeal electromyography (EMG) signal (Figure 3, A1 and A2, respectively; Supplementary Figure S 2) are smaller than those in response to i^2^CS with the opposite steering ratio (Figure 3, B1 and B2, respectively; Supplementary Figure S 2). Slow eCAP and respective breathing responses follow the opposite pattern (Figure 3, A3 vs. B3). To test the hypothesis that differential responses depend on interference rather than solely on activation of nearby vagal fibers by the two sinusoidal sources, sinusoidal stimulation with the same frequency and steering ratio was delivered through the same contacts, resulting in large, fast eCAP and EMG responses in both steering conditions (Figure 3, C1-2 and D1-2). On the other hand, breathing responses in the two steering conditions were similar to those with i^2^CS (Figure 3, C3, D3, see also Supplementary Figure S 3 for single contact stimulation).

**Figure 3:**
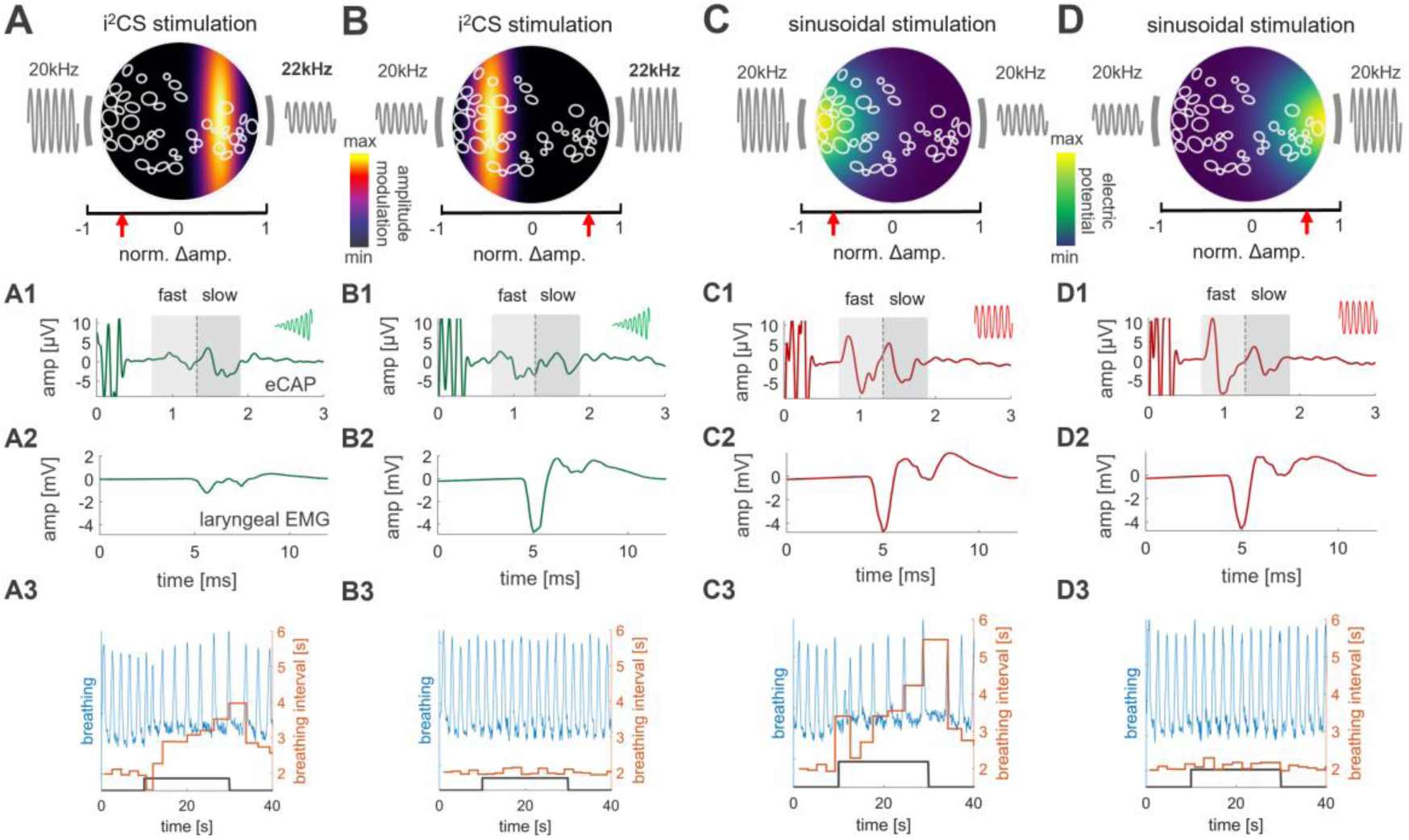
Intermittent interferential current stimulation (i^2^CS) elicits distinct experimental nerve and organ responses, which are different than those to equivalent, noninterfering sinusoidal current stimulation. (A) Schematic cross section of a stimulated VN at the level of an implanted MCE; shown are outlines of nerve fascicles and the 2 contacts (grey bars) used for i^2^CS, with the left source at greater amplitude than the right source (negative steering ratio expressed as the normalized difference in amplitude between both contacts; red arrow on left side of x-axis); left and right sources have carrier frequencies of 20 kHz and 22 kHz respectively. The colormap represents the maximum peak-to-peak amplitude of the beat interference envelope, indicating the location of the greatest modulation effect (cf. Supplementary Figure S 1). (A1) Evoked compound action potential (eCAP) response, triggered from the onset of i^2^CS, with 1.5 mA total current delivered through the 2 sources; slow and fast eCAP components are identified by the shaded areas corresponding to time windows defined by the average conduction velocities for ‘slow’ and ‘fast’ A-fibers. (A2) Weak laryngeal EMG response to i^2^CS stimulation. (A3) Large breathing response (blue) and respective change in breathing interval (orange) during a 20s-long train of i^2^CS stimuli (black trace). (B) Same as in A, but for the opposite steering direction (i.e., towards the right side). Sizeable eCAP and EMG responses, similar to left-steered stimulation, whereas breathing response is minimal. (C) Same as in A, but for sinusoidal stimulation. The two sources have the same carrier frequency (20 kHz). The strength of the electric potential generated by this particular current steering ratio is represented by a colormap. Strong, fast eCAP, EMG and intense breathing response. (D) Same as in C, but for the opposite steering direction. All eCAP and EMG responses are shown as averages of n = 660 trials.

*Experimental results suggest that interferential stimulation elicits specific nerve responses that depend on current steering and are different than those elicited by non-interfering, equivalent sinusoidal stimulation delivered through the same contacts*.

### 3. An anatomically realistic, physiologically validated model of the vagus nerve predicts that i^2^CS elicits reduced fiber activation at the focus of interference

To demonstrate interference at a focal area, responses of fibers within anatomically characterized organ-specific fascicles need to be recorded. Because recording from single fibers is not technically feasible, we used a recently developed modeling framework (Musselman et al., 2021) to compile an anatomically realistic neuro-electric model of a micro-CT-imaged and anatomically quantified swine vagus nerve (Figure 4, A-C). Because the particular nerve was stimulated in in vivo experiments, we were able to compare modeled and experimentally measured responses to the same interferential stimuli. The magnitude of the breathing response and the activity of modeled fibers in BP fascicles both change as a function of steering ratio, and are highly correlated (Figure 4, D); similarly, RL-mediated EMG responses and activity of modeled RL fibers are correlated (Figure 4, E). Significant correlations between measured physiological responses and modeled fiber responses are found across steering ratios (Figure 4, F), in line with significant correlations between experimental eCAP and physiological responses across steering ratios (Supplementary Figure S 2).

**Figure 4:**
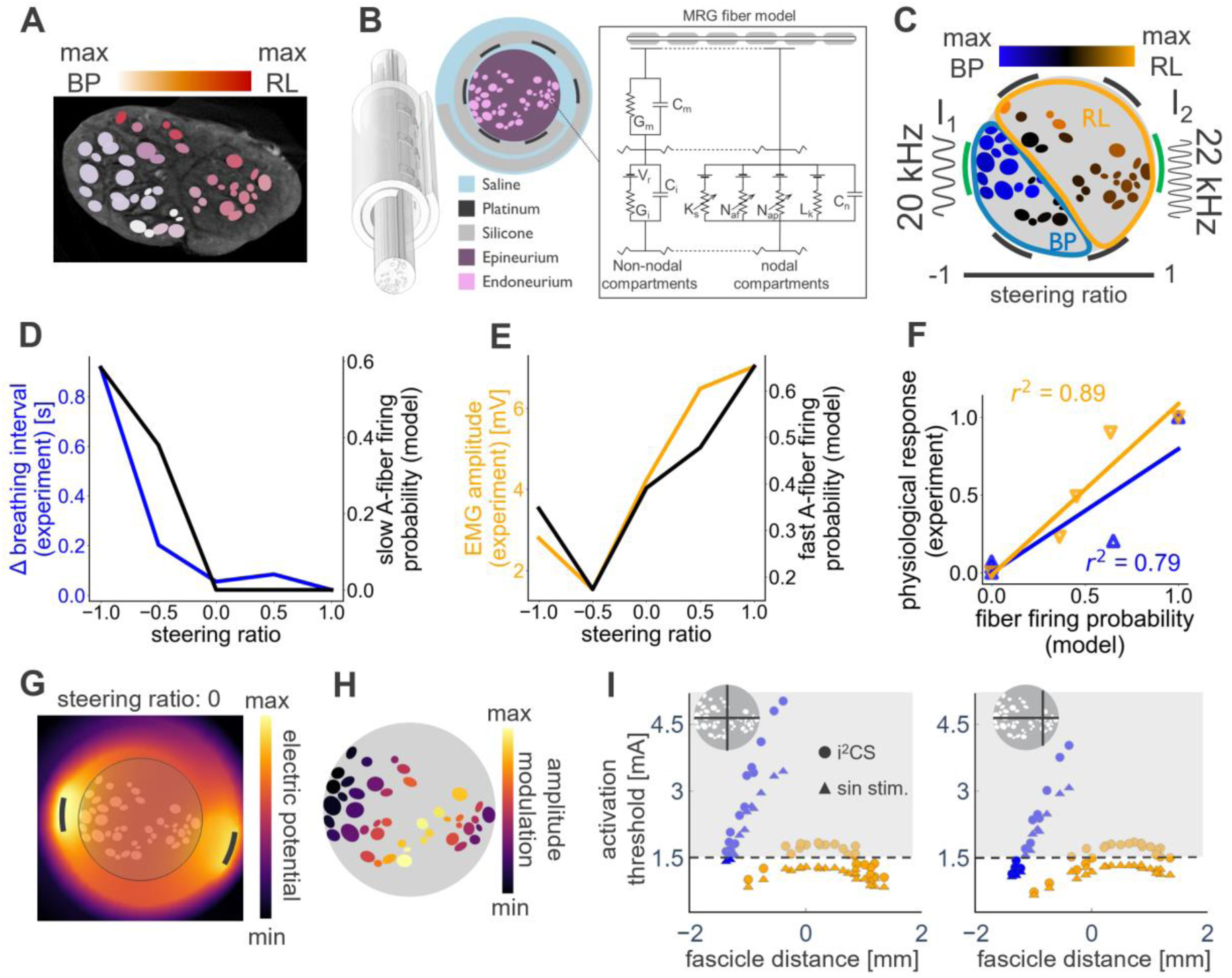
An anatomically realistic, physiologically validated biophysical model of the nerve-electrode interface predicts that i^2^CS produces reduced activation of fibers at the focus of interference. (A) Cross-section of micro-CT-imaged vagus nerve dissected from an experimental animal, at the level of the implanted cuff (same as in Figure 3). Fascicle color indicates the relative prevalence of BP (white) and RL fibers (red) within each fascicle. (B) Physical 3D model containing the nerve as an extrusion of the cross-section in (A) and the spiral cuff around it, including the different 3D domain materials and MRG-model (McIntyre et al., 2002) used to calculate the activation function of each fiber based on the electric field. (C) Cross-section of the nerve model after circular deformation, including the relative placement of (longitudinally positioned pairs of) contacts within the cuff (black lines); the circumferential position of 2 pairs of contacts used for stimulation are highlighted in green. One pair of contacts delivers a 20 kHz and the second a 22 kHz sinusoidal carrier. The horizontal (steering ratio) axis represents the ratio in stimulus amplitudes between the 2 contacts that controls the location of the interference focus. For visual clarity, areas with a predominance of RL (or BP) fibers are highlighted. (D) Modeled slow A-fiber responses to i^2^CS with different steering ratios (having combined total current amplitude of 1.5 mA) and change in breathing rate experimentally measured using i^2^CS with the same steering ratios. (E) Modeled fast A-fiber responses and EMG responses recorded experimentally. (F) Correlation between modeled normalized fiber firing probabilities and normalized physiological responses obtained experimentally in the same animal: fast A-fibers vs. EMG (orange), slow A-fiber vs. breathing response (blue). See Supplementary Figure S 5 for sinusoidal stimulation for panels D-F. (G) Map of the electric potential magnitude generated by i^2^CS with a total injected current of 1 mA and steering ratio of 0, focusing the amplitude modulation (AM) in the middle of steering axis. (H) Level of AM for all nerve fascicles under the same stimulation conditions in H. (I) Fiber activation threshold for i^2^CS (circles) and for equivalent sinusoidal stimulation (triangles) at BP (blue) and RL (orange) fascicles at different distances from the middle of steering axis for current steering towards the middle of the nerve (left) and towards the right (right). Insets indicate the focus of the interferential stimulation with a black cross, dotted black line indicates the current used for the in-vivo experiments and computational model (1.5 mA), and grey area indicates no activation.

Using the model, we estimated the magnitude of the interferential electric potential (Figure 4, G), the amplitude modulation (AM, Figure 4, H) and the activation thresholds of fibers (Figure 4, I, Supplementary Figure S 4), in different fascicles. Fiber activation thresholds inside fascicles experiencing maximum AM with i^2^CS are greater than with non-interferential sinusoidal stimulation, indicating reduced fiber activation at the focus of interference. In contrast, for fascicles closer to the nerve periphery, where non-interferential sinusoidal stimulation dominates, thresholds are similar for the two stimulation conditions (Figure 4, I).

*Results from anatomically realistic and physiologically validated biophysical models of vagal fibers indicate that i^2^CS results in increased fiber activation threshold at the focus of interference, compared to equivalent non-interferential stimulation, potentially providing an anatomical basis for selective VNS*.

### 4. Interferential stimulation activates vagal fibers in a specific spatiotemporal pattern in experimental recordings and in vagus nerve models

The time course of the amplitude modulation generated with i^2^CS depends on the difference between the two carrier frequencies, e.g., carrier frequencies of 20 and 22 kHz generate beats with 0.25 ms duration (Figure 1, D). In principle, the slower rise of AM at the focus of maximum interference results in gradual depolarization of fibers and a delay in the onset of i^2^CS-elicited responses, compared to the faster onset responses to sinusoidal stimulation (Figure 1, E). To test this hypothesis, we recorded laryngeal EMGs and eCAPs in response to i^2^CS and sinusoidal stimulation, at different steering ratios and beat durations (Figure 5, A; Supplementary Figure S 6). While sinusoidal stimulation elicits EMGs with the same short latencies independently of steering ratio, i^2^CS elicits EMGs with longer, steering ratio-dependent latencies (Figure 5, A), consistent with a shifting focus of interference. In addition, i^2^CS with different beat durations elicits EMGs with different latencies, all of which are longer than the latency of sinusoidal stimulation-elicited responses (Figure 5, B), consistent with slower activation of fibers by the rising AM at the focus of interference.

**Figure 5:**
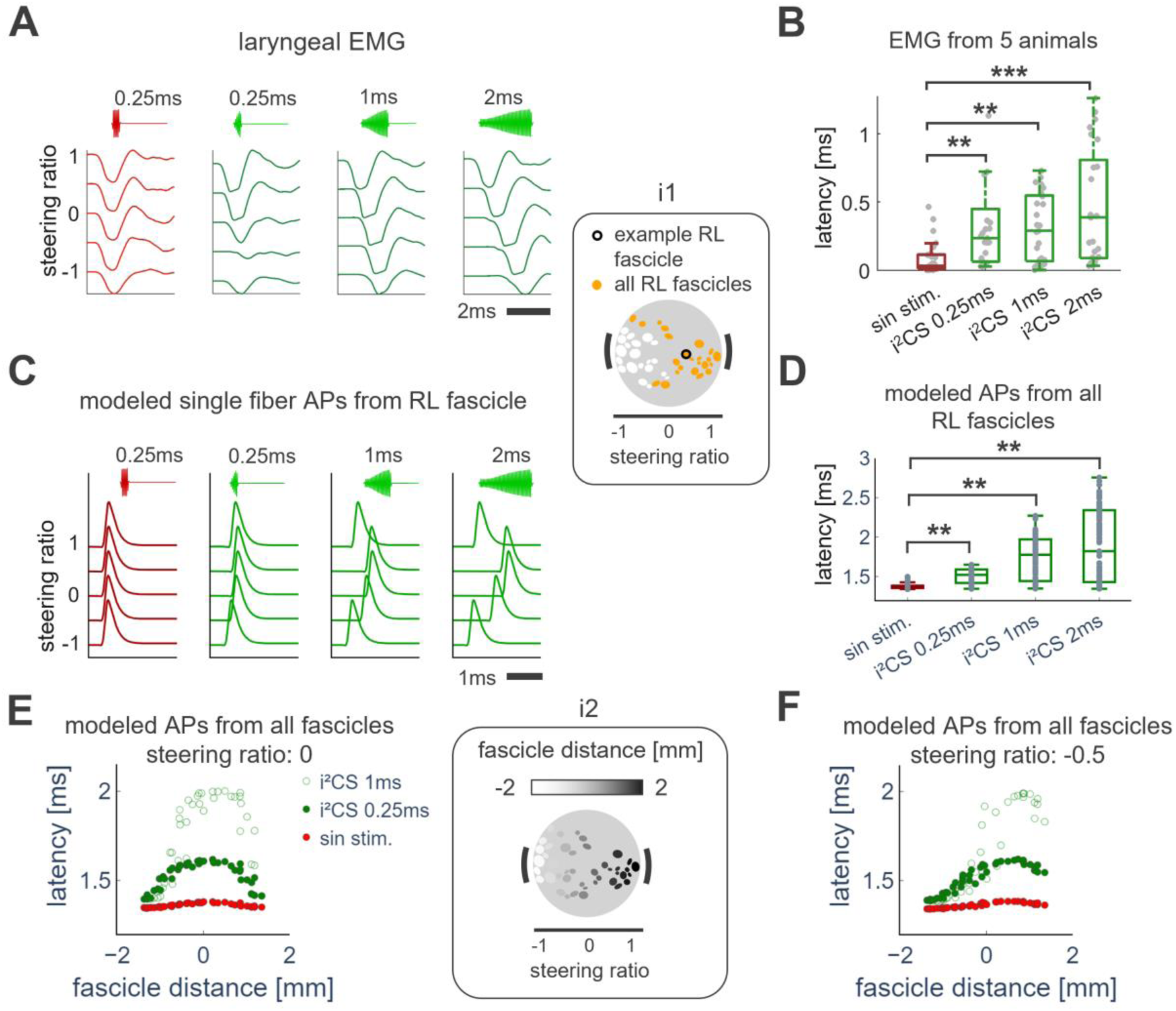
Interferential stimulation activates vagal fibers in a specific spatiotemporal pattern, in experiments and in vagus nerve models. (A) Laryngeal EMG responses from an example animal to a 0.25 ms long sinusoidal stimulus (red) and i^2^CS (green) at different steering ratios (total amplitude 2 mA) and beat durations (0.25, 1 and 2 ms, from left to right). (B) Difference in latency of onset of laryngeal EMG in response to 0.25 ms-long sinusoidal stimulation (red) and i^2^CS (green) of different beat durations (0.25, 1 and 2 ms, from left to right), across all steering ratios, from 5 animals. Response onset latencies to sinusoidal stimulation are shorter compared to i^2^CS of any beat duration (p<0.01, Kruskal-Wallis test). (C) Modeled APs in a fast A-fiber located in a deep, RL fascicle (black-outlined fascicle in inset i1), in response to sinusoidal (red) and i^2^CS (green), at different steering ratios (total amplitude 2 mA) and beat durations (0.25, 1 and 2 ms, from left to right), like those used in (A). (D) Difference in latency of onset of modeled APs calculated from simulations of fast A-fibers located in all RL fascicles (inset: orange-filled fascicles), for sinusoidal stimulation (red) and i^2^CS (green), at different beat durations (0.25, 1 and 2 ms, from left to right), across all steering ratios. AP latencies to sinusoidal stimulation are shorter compared to i^2^CS of any beat duration (p<0.01, Kruskal-Wallis test). (E) Onset latency of APs for modeled, fast A-fibers inside fascicles located at different distances from the mid-point of the steering axis, in response to sinusoidal stimulation (red data points) or i^2^CS with beat durations of 0.25 ms (filled green data points) and 1 ms (open green data points); the current was steered at the center of the nerve (steering ratio = 0; total amplitude 2 mA). Inset i2 shows modeled fascicles color-coded according to their distance from the mid-point of the steering axis. (F) Same as (E), but for a steering ratio of −0.5 (total amplitude 2 mA), resulting in an interferential field on the right side of the nerve cross-section.

To establish the single fiber basis of these effects, we modeled action potentials (APs) in response to i^2^CS in a deep RL fascicle (inset); we found that APs occur at different latencies depending on steering ratio, with slower onset of APs at fibers inside vs. outside of the interference focus (Figure 5, C); this finding is in agreement with experimental results obtained with i^2^CS-elicited eCAPs (Supplementary Figure S 6). Similarly, modeled APs elicited by i^2^CS with longer beat durations occur at longer latencies than those elicited by shorter beat i^2^CS or with sinusoidal stimulation (Figure 5, C), a difference that holds across all fascicles with a preponderance of RL fibers (Figure 5, D). In modeled fibers, increasing the beat duration of i^2^CS increases the latency of activation of fibers inside the focus of interference (Figure 5, E); the same dependency holds for a second interference focus, defined by a different steering ratio (Figure 5, F).

*Experimental and modeling results indicate that i^2^CS confers control of spatial and/or temporal aspects of activation of vagal fibers, by leveraging current steering and beat duration, respectively*.

### 5. Intermittent interferential stimulation controls precise timing of action potentials in modeled nerve fibers

Because interference produces a specific spatiotemporal pattern of fiber activation, the choice between continuous or intermitted stimulation may differentially impact generation of action potentials in nerve fibers. With i^2^CS, fascicles experience a range of AM levels, from minimal AM in superficial fascicles right next to contacts, to maximal AM in deeper fascicles (Figure 6, i1). Modeled responses to continuous iCS, with the same carrier frequency as i^2^CS used in the in vivo experiments (Figure 1, D), span a variety of profiles, depending on AM at the respective fascicle, e.g., phasic activation followed by block (Figure 6, A1, A2, and A5), regular tonic (Figure 6, A3) or irregular tonic activation (Figure 6, A4), in agreement with previous reports (Mirzakhalili et al., 2020). In contrast, intermittent i^2^CS with the same carrier frequencies and a pulse duration below the fibers’ refractory period (<2 ms), results in a predictable, regular temporal profile of fiber activation, across all fascicles regardless of their location, with an inter-spike-interval (ISI) determined by the pulse repetition frequency (33 Hz; Figure 6, B). Across all nerve fascicles experiencing different levels of AM (Figure 6, i2), the temporal precision of fiber activation, quantified by the variance of inter-spike interval (ISI) distributions, is relatively low for continuous iCS and depends on the location of the fascicle (Figure 6, C1-C3), whereas it is consistently high for intermittent i^2^CS, independently of the level of AM at the respective fascicle (Figure 6, C4).

**Figure 6:**
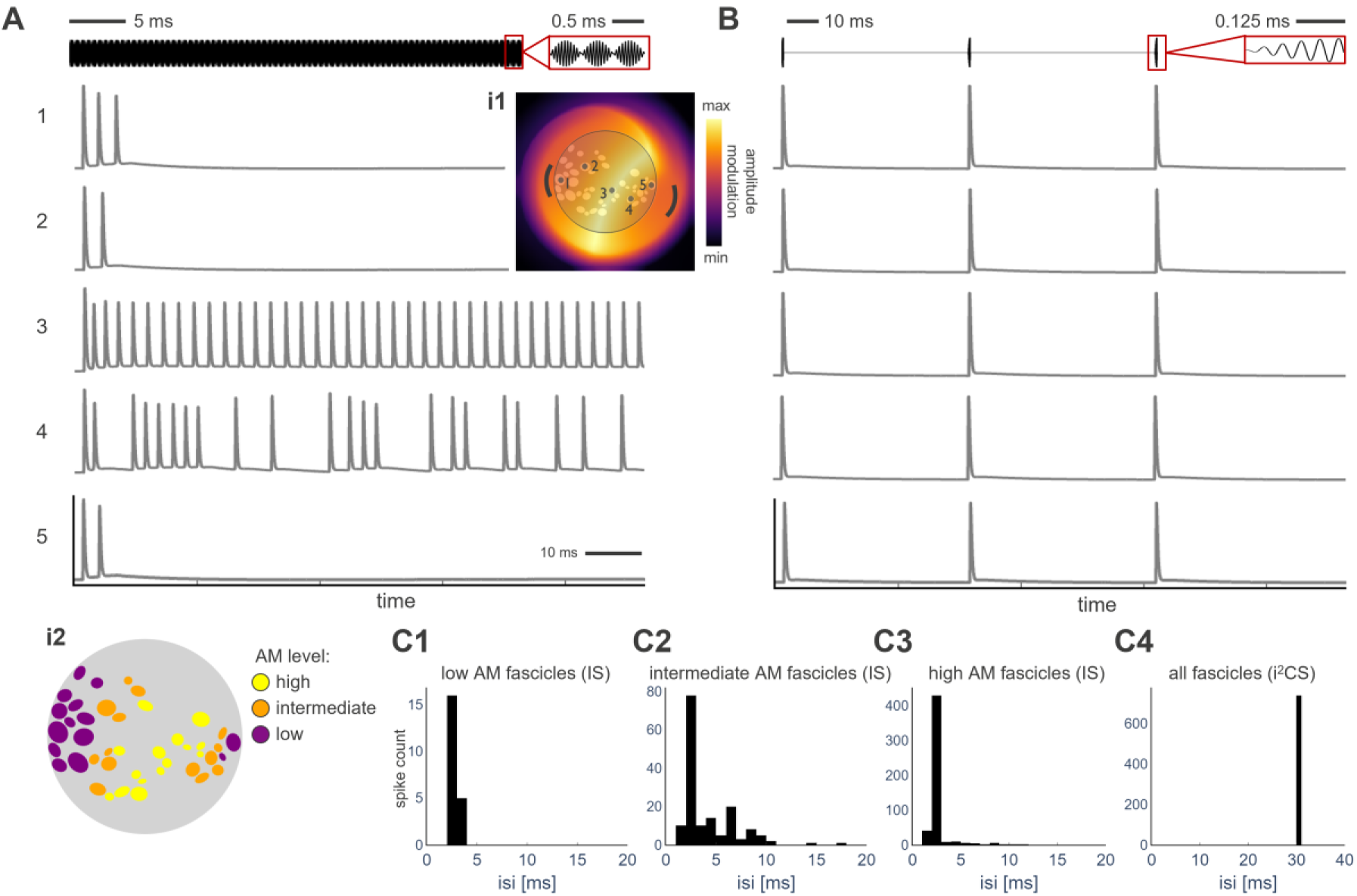
Repetition frequency of intermittent interferential stimulation controls timing of action potentials in nerve fibers in a temporally precise manner, in VNS models. (A) Modeled responses of fast A-fiber in several fascicles during continuous interferential stimulation. Stimuli with carrier frequencies of 20 kHz and 22 kHz and total amplitude of 2mA are deployed for up to 90 ms without interruption; stimulation signal at the top, with inset focusing on 3 consecutive beats. Traces 1-5 show the time course of responses of single fibers inside 5 fascicles, selected to demonstrate the effect of different levels of amplitude modulation (AM) of the electric field. Inset i1 shows the spatial distribution of the AM in a radial cross-section between cathodes and anodes of each source, where interference is strongest; numbers 1-5 indicate the selected fascicles. Fiber responses range from activation blocking (fascicles 1, 2, 5), to regular tonic firing (3), to irregular tonic firing (4). (B) Same as (A), but for i^2^CS, demonstrating regular firing in all 5 fascicles, with the inter-spike interval (ISI) being determined by the pulse repetition frequency (in this case 33 Hz, matching in vivo experiments). APs are elicited at different latencies across fascicles (cf. Figure 5, C-F, not visible here because of the long timebase). (C) ISI histograms obtained from APs from fibers in nerve fascicles exposed to different levels of AM (inset i2), for continuous interferential stimulation (IS): C1, fascicles with low AM (below first tertile), C2: intermediate AM (between first and second tertile), and C3: high AM (above second tertile). C4: for i^2^CS (all fascicles).

*Neural modeling results indicate that intermittent interferential stimulation precisely controls the timing of elicited action potentials in fibers across the entire nerve, with ISIs determined by the pulse repetition frequency*.

### 6. Interferential stimulation has improved functional selectivity for a desired effect over a side effect compared to equivalent, non-interfering sinusoidal stimulation

The spatial distributions of RL and BP fibers along a transverse axis show separate peaks at deep-lying fascicles rather than at the nerve periphery (Figure 2, D5). We therefore hypothesized that interferential stimulation producing maximum AM in deeper fascicles on the “RL side” of the nerve would result in reduced activation of an RL-mediated side-effect (laryngeal muscle contraction) over a BP-mediated desired effect (breathing response), compared to equivalent, non-interfering sinusoidal current stimulation. We recorded nerve potentials (eCAPs) in response to i^2^CS and to sinusoidal stimulation, at different steering ratios; we found that, while slow eCAP responses, corresponding to the smaller A-fiber-mediated, desired effect, were similar in both conditions, i^2^CS elicited smaller fast eCAPs, which correspond to the larger A-fiber-mediated side effect (Figure 7, A and B, respectively). Compared to sinusoidal stimulation, i^2^CS is associated with both higher selectivity and greater range of slow eCAPs across several steering ratios (Figure 7, C), resulting in greater selectivity for smaller A-fibers in several animals (Figure 7, D). Similarly, i^2^CS produces the same level of the breathing response (Figure 7, E), but with smaller amplitude of laryngeal EMG (Figure 7, F), resulting in greater selectivity for the desired effect, both in individual animals (Figure 7, G) and across several animals (Figure 7, H). These findings indicate that by adjusting the steering ratio of i^2^CS, lung- and larynx-specific responses are shifting according to the idea of reduced fiber activation, which is consistent with the bimodal anatomical distribution in the vagal trunk (Figure 2, Supplementary Figure S 7). Additionally, i^2^CS attains improved selectivity compared to square pulse VNS delivered through a multi-contact cuff electrode (Supplementary Figure S 8). Likewise, in our computational model of the vagus nerve, firing probabilities of smaller fibers inside BP fascicles and of larger fibers inside RL fascicles have steering ratio-dependent activation profiles consistent with experimental measurements, for i^2^CS and sinusoidal stimulation (Figure 7, I-K), which result in higher selectivity for BP fibers with i^2^CS (Figure 7, L).

**Figure 7:**
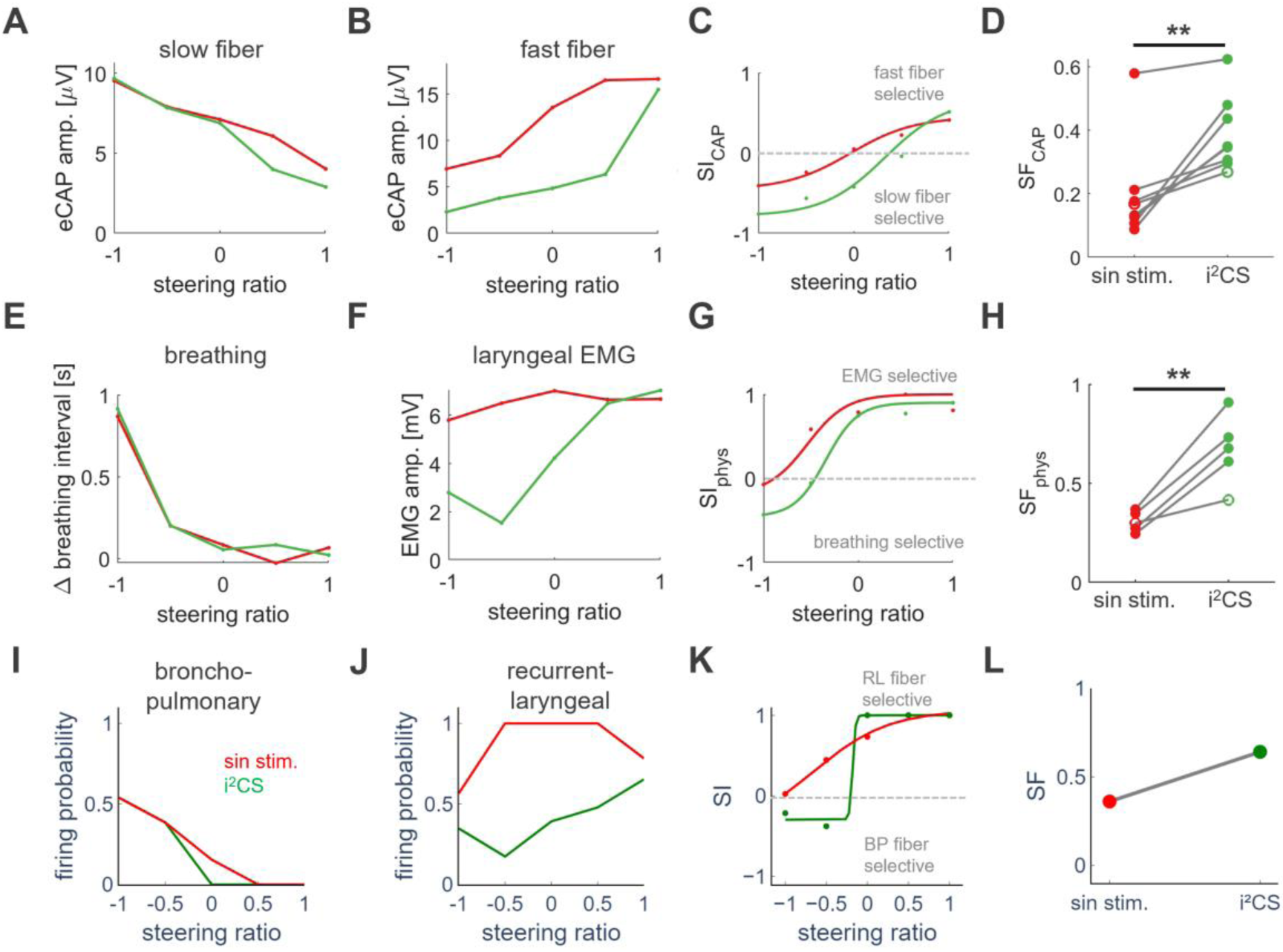
Interferential stimulation attains increased selectivity of a desired effect, mediated by smaller BP fibers, over a side effect, mediated by larger RL fibers, compared to equivalent sinusoidal stimulation. (A) Slow eCAP amplitudes for sinusoidal and interferential stimulation at different steering ratios, from an example animal. (B) Same as in (A), but for fast eCAPs. (C) Slow over fast eCAP selectivity index (SI) defined as the ratio of the eCAP amplitudes in A and B, fitted with a sigmoidal function, for the 2 stimulus conditions. (D) eCAP selectivity factor (SF), defined as the product of the slope and range of the fitted sigmoidal function of the SI (Supplementary Figure S 9) for the 2 stimulus conditions in 8 animals (example animal denoted with open symbols) (p = 0.007; Wilcoxon rank-sum test). (E) Magnitude of the (desired) breathing response (change in breathing interval, ΔBI) at different steering ratios, from an example animal, for interferential and equivalent sinusoidal stimulation. (F) Amplitude of the (undesired) laryngeal EMG at different steering ratios, in the same animal. (G) Physiological SI, defined as the ratio of the magnitude of the desired over the side effect, in the same animal. (H) Physiological SF, comparing interferential and equivalent sinusoidal stimulation in 5 animals (open symbols: example animal) (p = 0.008, Wilcoxon rank-sum test). (I) Firing probability of BP fibers (modelled as smaller A-fibers, diameter 6 μm, placed inside fascicles rich in BP fibers) for sinusoidal (red) and interferential (green) stimulation at different steering ratios and a total stimulation amplitude of 1.5 mA. Results obtained using the anatomically realistic biophysical model of the example animal. (J) Same as in A, but for RL fibers (modelled as larger A-fibers, diameter 10 μm, inside fascicles rich in RL fibers). (K) BP over RL SI calculated from the firing probabilities in A and B, fitted with a sigmoidal function. (L) SF comparing the sinusoidal and interferential stimulation conditions.

*In a series of experiments in swine, i^2^CS having maximum interference focus on RL fascicles has improved selectivity for a desired effect, mediated by smaller BP fibers, over a side-effect, mediated by larger RL fibers, compared to equivalent non-interfering sinusoidal stimulation*.

## Discussion

### I^2^CS activates vagal fibers in a spatially focused and temporally precise manner

Our report demonstrates that interferential stimulation is a viable method for tunable and precise, spatially selective VNS. Selection of the 2 MCE contact pairs for i^2^CS defines the steering axis on the radial plane, and of the 2 stimulus intensities (steering ratio) defines the maximum AM site along that axis (Figure 3, Figure 4, Figure 5). To the best of our knowledge, ours is the first demonstration, both in principle and in practice, of increased organ selectivity due to the improved control of the spatial focus at which the maximum AM of the electric field is generated. This is important because, to the extent that the vagus nerve in humans has an organotopic fascicular organization (Jayaprakash et al., 2023; Kronsteiner et al., 2024), spatial focusing may provide a strategy for selective VNS (Ahmed et al., 2022). Selective VNS may minimize undesired responses from non-targeted organs, thereby improving dose titration and therapeutic efficacy (Gorman and Mortimer, 1983). For example, even though VNS in epilepsy is generally safe and well tolerated in the long-run, titration of therapy is performed progressively, over repeated office visits, to minimize side effects like cough and voice alteration, arising from activation of large, low threshold laryngeal and pharyngeal fibers (Heck et al., 2002); rapid titration could significantly accelerate clinical response, as reported in a recent meta-analysis (Tzadok et al., 2022). Similarly, in clinical studies of VNS in heart failure, laryngeal and pharyngeal side effects prevented clinicians from adequately dosing VNS to a level required to activate smaller, higher threshold cardiac fibers mediating the desired effect of cardio-inhibition, possibly contributing to the failure of clinical trials (Gold et al., 2016).

Our study also demonstrates that intermittent delivery of short ‘pulses’ of interfering stimuli results in temporally precise activation of vagal fibers, with the timing of elicited APs controlled by the amplitudes and frequencies of the 2 interfering sources (Figure 5), and ISIs controlled by the pulse repetition frequency (Figure 6). Standard, square pulse VNS elicits temporally precise, action potentials time-locked to the stimulus, but its spatial selectivity is limited. On the other hand, suprathreshold (Chang et al., 2022; Pelot et al., 2017) or subthreshold high-frequency stimulation (Vargas et al., 2023) attains improved fiber selectivity but elicits asynchronous action potentials in nerve fibers, with limited precision. To the best of our knowledge, i^2^CS is the first stimulation paradigm that combines spatial focusing with temporal precision. Temporally precise stimulation of vagal fibers is useful when fiber activation needs to be tightly controlled relative to a dynamically changing physiological state. For example, respiratory-gated auricular nerve stimulation is thought to control hypertension by eliciting afferent volleys at specific phases of the respiratory cycle, when sensory brainstem neurons involved in the baroreflex are more excitable (Garcia et al., 2017; Sclocco et al., 2017). Likewise, delivering vagal stimuli at specific phase of the cardiac pacemaker cells during the cardiac cycle may differentially impact the risk of vagally-induced sinus, atrial, sinoatrial or ventricular arrhythmias (Goto et al., 1983; Jalife et al., 1983; Jalife and Moe, 1979; Kharbanda et al., 2022; Slenter et al., 1984). Finally, closed-loop VNS to control blood pressure (S. Zanos, 2019), treat arrhythmias (Ottaviani et al., 2022), modulate gastric sphincter function in gastrointestinal disorders (Payne et al., 2019) or regulate inflammation-related functions of the vagus nerve (T. P. Zanos, 2019) relies on delivering precise, short latency responsive stimulation after a relevant physiological event is detected, a scenario feasible with the use of i^2^CS. Recent reports have demonstrated the feasibility of dedicated miniaturized and low-power integrated circuits capable of delivering iCS to peripheral nerves (H. Xin et al., 2024) and of methods to efficiently capture and read out neural responses to stimulation (Y. He et al., 2022). Due to its intermittent charge delivery method, power consumption of i^2^CS is similar to standard biphasic current stimulators, i.e., i^2^CS is much more power efficient than its continuous counterpart paving the way for VNS devices capable of long-term closed-loop, spatio-temporal control of fiber activation.

### Sources of selectivity and mechanisms of action of i^2^CS

Counterintuitively to the expectation that activation of fibers will be more efficient in the focal point of interference, our experimental and modeling results indicate that i^2^CS achieves greater selectivity for a desired bronchopulmonary (BP) response over an undesired recurrent laryngeal (RL) response, compared to sinusoidal stimulation (Figure 7, E-H) or to square pulse VNS (Supplementary Figure S 8), in a different manner: Surprisingly, improved selectivity with i^2^CS is driven mostly by reduced laryngeal EMG for a given level of breathing response, when maximum AM is focused on RL fascicles, thereby eliciting reduced RL fiber activation (i.e., negative steering ratios; Figure 3; Figure 7, E-H). In contrast, noninterfering sinusoidal stimulation produces less of a graded laryngeal EMG response along the steering axis (Figure 7, E-H), as fibers in RL fascicles, in the absence of AM, are consistently activated by the sinusoidal currents (Figure 7, I-L). Consequently, slow eCAPs, generated by smaller A-fibers, some of which innervate the lung, are preferentially elicited over fast eCAPs, generated by larger fibers, some of which innervate the larynx (Figure 7, A-D), and firing probability of smaller fibers in BP fascicles is greater than for larger fibers in RL fibers, when the field is focused on BP-rich fascicles (e.g., Figure 7, I vs. J, for negative steering ratios). In addition to reduced side effect, improved selectivity may permit testing of a larger range of stimulus intensities for calibration of the desired effect (Figure 7).

The difference in selectivity between i^2^CS and sinusoidal stimulation likely arises because fibers show lower activation threshold when exposed to a non-amplitude modulated sinusoidal (20 kHz – 22kHz) field compared to an amplitude-modulated field (2kHz beat frequency) (Figure 4, I, Supplementary Figure S 10). Amplitude-modulated fields with progressively increasing charge per depolarization-hyperpolarization cycle likely result in slower net depolarization, which has been linked to reduced fiber activation for a given intensity level (Hennings et al., 2005; Vuckovic et al., 2008). Also, the charge per cycle in the case of 2 signals of equal carrier frequency (sinusoidal stimulation) is greater than when one of the 2 carrier frequencies is higher than the other, as in the case of interferential stimulation. Moreover, the higher that carrier frequency, which results in shorter beat duration, the lower the charge per phase for the amplitude modulated signal.

Importantly, our experimental and modeling results provide evidence that the anatomical substrate, i.e., organ-specific fibers, upon which spatially selective stimuli are applied explains much of the variability in the physiological responses, i.e., organ-specific effects (Figure 4). This underlines the practical significance of resolving the functional anatomy of nerves and using anatomical constraints in the design of nerve interfaces (Musselman et al., 2023). In our experiments in swine, almost no electrode pair or steering ratio was associated with perfect selectivity for the desired, BP-mediated breathing effect (Figure 7, D, H). This is likely due to the significant mixing of BP and RL fibers inside the same fascicles (Figure 2, D2-D4), which poses fundamental anatomical limitations in the degree of functional selectivity of any stimulus targeting single fascicles or small groups of fascicles. Attaining greater selectivity would require sub-fascicular stimulus resolution, e.g., by using more than 2 current sources for interference, or by using high-channel count intraneural electrode arrays that can target smaller sub-fascicular sectors or even single fibers (Badi et al., 2021).

### Study limitations

Our study has several limitations. First, our methodology for anatomical tracking cannot reconstruct trajectories of single fibers and assumes that mixing of fibers when 2 fascicles merge into one is uniform across the resulting fascicle (Figure 2). This assumption does not consider sub-fascicular organization of fibers (Jayaprakash et al., 2023) and may result in an overestimate of the amount of fiber mixing at the cervical level. Second, although our models are anatomically realistic and experimentally validated (Figure 4, Figure 5), they are not ideal. For example, the nerve in our model is deformed to a circular shape and fascicles are modeled as extrusions of a single nerve cross section instead of more complex splitting and merging structures, thereby limiting accurate modeling of electrical fields (Ciotti et al., 2024). Fiber populations linked to desired and side effects are simplified by modeling a single fiber inside each fascicle with one of two sizes and simplified ionic conductances, instead of modeling many fibers, with a variety of sizes, specific sub-fascicular clustering statistics and a variety of ionic conductances (Ciotti et al., 2024; Jayaprakash et al., 2023; Pelot et al., 2021). Third, our experimental and modeling approaches do not consider current shunting and escape of current outside of the cuff, both of which are likely altering physiological responses significantly (Blanz et al., 2023; Nicolai et al., 2020). Finally, we did not study i^2^CS delivered through chronically implanted cuffs or in awake animals; stimulation responses in both these cases are likely to be different than those in acutely implanted, anesthetized animals reported here, as shown previously (e.g., Ahmed et al., 2021).

### Conclusions

In this work we have introduced a new electrical stimulation paradigm, called intermittent interferential current stimulation (i^2^CS), that allows for tunable and precise spatiotemporal control of fiber activation during PNS. As a result, i^2^CS demonstrates improved selectivity for a desired effect over a side effect, when compared with standard sinusoidal or square pulse stimulation. We have also uncovered a new mechanism of action of i^2^CS, which includes reduced and delayed fiber activation at the focus of interference. Compared with previously proposed, continuous interferential methods, i^2^CS is more energy efficient and can be readily implemented in standard implantable stimulation devices.

## Acknowledgments

This work was partially supported by NIH SPARC 75N98022C00019 to SZ. IMEC has been granted patents related to this work (US18511762, US0017074A1, US0364428A1, US0198109A1). IMEC and Northwell have submitted patents related to this work. The authors wish to thank Patrick van Deursen and Yousef Al-Abed for supporting the collaboration between IMEC and the Feinstein Institutes for Medical Research, Evelien Hermeling for the support on data analysis, and Eva Severijnen for the support on the design of in vivo experimental protocols.

## Materials and methods

### Animals and surgery

The experimental protocol used in this study has been described in detail earlier (Jayaprakash et al., 2023). In brief, the effects of i^2^CS on physiological and neural response were examined in 8 male Yucatan swine (30-54kg). All animal protocols and surgical procedures were reviewed and approved by the animal care and use committee of Feinstein Institutes for Medical Research and New York Medical College. Animals were sedated with a mixture of Ketamine (10-20 mg/kg) and Xylazine (2.2 mg/kg) or Telazol (2-4 mg/kg). Propofol (4-6 mg/kg, i.v.) was used to induce anesthesia, and following intubation, the anesthesia was maintained with isoflurane (1.5-3%, ventilation). Body temperature was maintained at 38-39°C using a heated blanket. Blood pressure and blood oxygen level were monitored with a cuff and a pulse oximeter sensor. All surgeries were performed using sterile techniques.

### Cervical vagus and laryngeal muscle implants

The cervical vagus nerve was exposed and 2 multi-contact cuff electrodes (MCEs, Cortec GmbH) were placed on the nerve. The MCE for stimulation (custom AirRay spiral, 18 contacts, see Supplementary Figure S 11) was placed rostrally, ∼2 cm away from the nodose ganglion. A second recording MCE (AirRay helix cuff) was placed 5-6 cm caudally from the stimulation MCE to record eCAP waveforms. Electrode impedances at a frequency of 1 kHz were measured in vivo using an IMP-2A impedance tester (Microprobe) to verify good contact with the tissue. For laryngeal muscle recordings, Teflon-insulated single or multi-stranded stainless-steel wires were de-insulated at the tip for about 1 mm and inserted in the thyroarytenoid laryngeal muscle with a needle. In 3/8 animals, the laryngeal EMG signal deteriorated and was lost over the course of the experiment, preventing the calculation of physiological selectivity indices (cf. Figure 5 H, L).

### Experimental setup

The experimental setup as described in Supplementary Figure S 12 was deployed for in-vivo VNS. Stimulation waveforms and digital signals for timing pulses and stimulation trains were designed using Python 3.9 (Van Rossum and Drake, 2009) using a sampling frequency of 1 MHz and transmitted from a PC to a data acquisition (DAQ) board (NI-PCIe6363, National Instruments) via serial communication. The parameters of the two types of stimulation waveforms (sinusoidal and i^2^CS) are listed in Table 1. A pulse repetition frequency of 33 Hz was chosen to avoid noise at harmonic multiples of the power line frequency (60 Hz), and low enough to avoid muscle fatigue (cf. Supplementary Figure S 13). Stimulus presentation was randomized.

**Table 1:** Stimulation waveforms for in vivo experiments.

| Waveform type | Burst duration [ms] | First carrier frequency [kHz] | Second carrier frequency [kHz] | Pulse repetition frequency [Hz] | Train duration [s] |
| --- | --- | --- | --- | --- | --- |
| Sinusoidal | 0.25 | 20 | 20 | 33 | 20 |
| i<sup>2</sup>CS 0.25ms | 0.25 | 20 | 22 | 33 | 20 |
| i<sup>2</sup>CS 1ms | 1 | 20 | 20.5 | 33 | 20 |
| i<sup>2</sup>CS 2ms | 2 | 20 | 20.25 | 33 | 20 |

The stimulation waveforms were used as input for the DAQ to generate analog output waveforms, and voltage to current conversion was performed via custom-made dual differential Howland current pumps with 1 V : 10 mA conversion factor and power supply of +15/−15 V. To ensure that the stimulation sources were isolated from the rest of the hardware, the Howland current pumps were powered using a 22.5W, 20000mAh battery power bank (INIU) and the two outputs of the Howland current source were connected to the stimulation cuff via a custom analog multiplexer designed to allow each independent current source to be routed to any of the electrode channels. The channel selection was controlled digitally using the DAQ system. The connection from the multiplexer to the spiral stimulation cuff was implemented via a micro 360 plastic circular straight tail connector (Omnetics Connector Corporation).

Two additional digital signals were generated by the DAQ: one pulsed digital line whose value was set to 5V during each stimulation burst and −5V otherwise, and a stimulation train line whose value was set to 5V during the whole duration of the stimulation train and 0 otherwise. The two digital lines were directly connected to the digital input ports of the recording instrumentation for synchronization during data acquisition.

### Measurement of physiological and neural signals

All physiological signals were continuously sampled at 1 kHz (PowerLab 16/35, ADI) and visualized using LabChart (ADI). We monitored heart rate by recording ECG in a 3-lead patch electrode configuration from the wrist of the animal. Signals were amplified using a commercial bio-amplifier (FE238, ADI). Breathing rate was monitored via a respiratory belt transducer (TN1132/ST) connected to a bridge amplifier (FE221, ADI). The train digital line from the DAQ was used to identify the stimulation windows for post-processing.

Neural and EMG signals were sampled at 30 kHz using a second data acquisition system including an amplification and digitization head-stage and controller unit (RHS-32, Intan Tech). The train digital line from the DAQ was used to trigger the start and stop of the recording, and the pulse digital signal was used to identify single burst stimulation windows for post-processing.

### Analysis of physiological and neural signals

Raw physiological signals were high-pass filtered post-hoc to remove the DC-component (4-pole Butterworth filter, cut-off frequency: 0.1 Hz) and a custom-made beat-detection algorithm was used to extract the heart rate (HR) and the breathing interval (BI). The increase in BI was used as a measure for stimulation effectiveness since stronger stimulation of the vagus nerve can sometimes lead to apnea, which cannot be quantified via breathing rate reduction. Response strength was calculated as the average BI in the stimulation window (20 s) corrected by the average baseline BI in a 10 s window before stimulation onset. Interstimulus interval was 60 s.

Raw neural signals were subjected to a 60Hz notch-filter implemented either with the Intan recording software or post-hoc in MATLAB. Furthermore, signals were also high-pass filtered post-hoc (4-pole Butterworth filter, cut-off frequencies: 10 and 260 Hz for EMG and eCAP, respectively) before averaging the responses over all stimulus presentations in the train (default: 33 Hz inter-pulse-interval (IPI) for 20 s, i.e., 660 pulses). EMG response strength was calculated as the peak-to-peak magnitude of the signal in a pre-defined response window after stimulation (4-12 ms). eCAP response strengths were calculated in 2 pre-defined response windows after stimulation onset that were derived from conduction velocities of ‘fast’ and ‘slow’ A-fibers and the measured distance between the stimulation and recording electrodes.

To quantify the effect of stimulation parameters on fiber and organ functional selectivity, we defined selectivity indexes as previously described (Chang et al., 2022), and adapted to account for the different experimental conditions. The selectivity index *SI* was calculated for eCAP (1) and physiological (2) responses as follows:

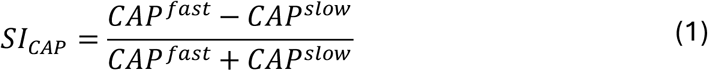

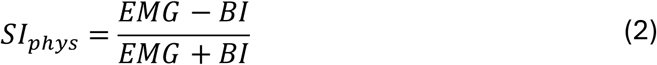

To quantify the dependence of selectivity on steering, the data was fitted with a modified logistic function of the form described in (3):

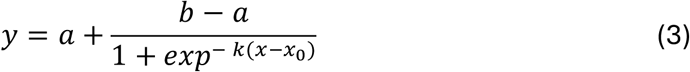

Where a, b, k and x_0_ are the minimum and maximum reachable values of the fit, the steepness of the slope and the inflection point, respectively. The Selectivity Factor *SF* was derived from the logistic fits to the SI-data as a product of 2 parameters as described in (4), where 〈*slope*〉 and 〈*range*〉 are the normalized slope and range of the SI sigmoidal function in (3):

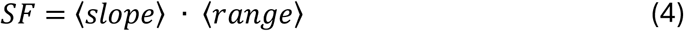

The normalized slope is defined as the sigmoid slope converted to degrees and normalized by the maximum value of 90° (Supplementary Figure S 9A). Here, the slope value stands for the ‘cutoff sharpness’ of the change of selectivity across the nerve diameter while the range describes the maximum relative difference between the activation of one function versus the other. The values for range and slope are reported separately in Supplementary Figure S 9B.

All data analysis was performed using custom-made or publicly available scripts in MATLAB (Mathworks).

### Quantification of nerve anatomy

#### Anatomical dissection and micro-CT imaging of nerve samples

Animals were euthanized by injection of Euthasol (1 ml/10 pounds BW, i.v.); death was confirmed using ECG and absence of arterial pulse. After euthanasia, cervical vagus nerves were dissected from above the nodose ganglion to the end of the thoracic vagus; during that time, nerve branches, still attached to the nerve trunk, were isolated using blunt dissection up to the respective end organ (heart, lung or larynx). A fine suture loop (6-0) was placed on the epineurium of each branch, close to its emergence from the trunk, to label that branch and maintain a record of the innervated organ during subsequent imaging studies, as the sutures used are radiopaque and visible in micro-CT images. The nerve trunk, along with organ-specific labels was photographed before and after extraction. The samples were fixed in 10% formalin for 24 h, then transferred to Lugol’s solution (Sigma, L6146) for five days to achieve optimal fascicular contrast for the micro-CT scan. Nerve trunks were sectioned into several 6 cm-long segments and the rostral end of each segment was marked with a suture knot, to maintain the rostral-caudal direction. Each nerve segment was scanned individually in the micro-CT scanner. Nerve segments were mounted in position on a vertical sample holder tube. The samples were scanned using Bruker micro-CT Skyscan 1172 with a voxel size of 6 μm using.

#### Fascicle tracking

After the following parameters: 55 kV, 149 μA, 0.5mm Al filter, rotation step of 0.5, and frame averaging of 6. During reconstruction of the images using cone-beam reconstruction software based on the Feldkamp algorithm (Skyscan NRecon, version 2.2.06), a ring artifact correction of 5, a beam hardening correction of 40%, was applied to all samples, as was automatic post-alignment. Reconstructed cross-sectional image slices were saved as bmp files.

#### Fiber mixing model

Each node in the graph was also assigned a tuple representing the percentage of the main trunk, the recurrent laryngeal branch, and the bronchopulmonary branch. These structures were manually identified and the leaf nodes in the graph corresponding to these structures were assigned to 100% for the respective structure and 0% of the other structures. The percentages were propagated through the graph from the caudal to the cranial end based on the following rules:

a. If a node has only a single input connection from the previous node, then the percentages do not change. This is true even if the previous node branches into multiple nodes. This method assumes homogenous mixing.
b. If a node contains multiple input connections from multiple previous nodes, as is the case when merging, then the percentages in the next node is the weighted average of the percentages in the previous nodes with a normalized weighting based on the areas of the previous ellipses. This method assumes instantaneous mixing at merge locations.

#### Spatial distribution of fibers

The spatial distributions of RL and BP fibers was determined by projecting the centroids of each fascicle onto a line passing through the center of mass for each fiber type. The mass of each fiber type for a given fascicle is proportional to the area of the fascicle times the percentage of each fiber type, assuming a constant fiber density. Histograms of values proportional to the fiber counts for each fiber type were generated along the axis of the line.

#### Computational Models

Anatomically realistic 3D computational models were obtained via the adaptation of the ASCENT framework (Musselman et al., 2021). In this study we deployed the ASCENT framework using Python 3.9, COMSOL Multiphysics 5.6 (COMSOL Inc., Burlington, MA), and NEURON v7.6 (Hines and Carnevale, 1997). The framework allows to construct a 3D nerve anatomy starting from an anatomical image and extruding the nerve section over the desired length of the nerve fiber. For this purpose, a nerve cross section from a location right below the middle row of electrodes in the nerve cuff placed on an example animal was selected to generate the 3D model (Supplementary Figure S 11). The nerve shape was deformed to account for circular deformation after cuff placement ensuring a minimum inter-fascicle distance and fascicle distance to the nerve perimeter of 10 um. A scale ratio µm/pixel of 1.56 was used to account for histological tissue shrinkage and to reach a final nerve diameter of ≈ 3mm as measured experimentally during the experiment. The perineurium was modelled using a thin layer approximation whose parameters were derived from pig Vagus nerve experiments (Pelot et al., 2020), and the perineurium conductivity was selected to be dependent on temperature and frequency, with values set at 37 °C and 20 kHz to account for the stimulation frequency (Weerasuriya et al., 1984). Each fascicle was populated with a single myelinated fiber placed in the middle and modelled using the MRG model (McIntyre et al., 2004, 2002) using a fiber diameter as detailed in the analysis.

A 3D model of the electrode cuff placed around the nerve was obtained starting from the electrode geometrical specifications (Supplementary Figure S 11). Material properties were assigned to the different electrode components as follows: electrode contacts (Pt), cuff insulation (silicone), and fill medium and recess (saline). The proximal and outer medium were defined as cylinders with radius 3.5 mm and 4.3 mm respectively.

The voltage transient and activation threshold simulation protocols in ASCENT were used to derive fiber responses resulting from electrical stimulation. As the ASCENT pipeline does not allow to simulate multiple waveforms at the same time as required during i^2^CS, the framework was extended to support this use case. The extension introduces the summation of the electric potentials resulting from separate simulation waveforms. The electric potential over the 3D space resulting from the injection of a DC current at the electrode sites was computed, and then modulated by the stimulation waveform, which allows to obtain a temporal profile of the electric potential during stimulation. In the present work, we computed separately the modulated electric potential for each one of the current sources, and then leveraged the principle of superposition of effects to sum the two electric potential temporal profiles, which allowed to obtain a single temporal profile of the electric potential resulting from the presence of two separate current sources. The resulting potential was then used in the pipeline to derive fiber responses. We simulated different stimulation waveforms, replicating the experimental conditions as described in Table 2.

**Table 2:** Simulation parameters.

| Waveform type | Burst duration [ms] | First carrier frequency [kHz] | Second carrier frequency [kHz] | Pulse repetition frequency [Hz] | Number of repetitions |
| --- | --- | --- | --- | --- | --- |
| Sinusoidal | 0.25 | 20 | 20 | 33 | 3 |
| i<sup>2</sup>CS 0.25ms | 0.25 | 20 | 22 | 33 | 3 |
| i<sup>2</sup>CS 1ms | 1 | 20 | 20.5 | 33 | 3 |
| i<sup>2</sup>CS 2ms | 2 | 20 | 20.25 | 33 | 3 |

In the case of continuous interferential stimulation (IS), the same carrier frequency used for i^2^CS 0.25 ms (20 kHz and 22kHz) were used, each carrier having a total duration of 90 ms, which is equivalent to the total duration of the i^2^CS 0.25 ms stimulus train used for comparison.

For each waveform, current amplitudes of 1.5 mA and 2 mA were used as they replicate current amplitudes used during experiments. Moreover, for each current amplitude, five different amplitude ratios between current sources were simulated, as defined in (5): 0.9, 0.7, 0.5, 0.3 and 0.1.

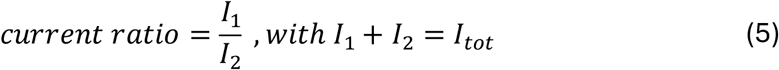

The steering values were mapped to a −1 to +1 range for illustration purposes (i.e., 0.9 = −1, 0.7 = −0.5, 0.5 = 0, 0.3 = 0.5, 0.1 = 1).

#### Analysis of electric potential

Electric potential maps were generated from COMSOL injecting a 1 mA DC current at each contact electrode and weighting the resulting electric potential by the contribution of the single source for the specific steering ratio (for example a factor of 0.5 is used for both sources in the case of steering ratio 0). The two potentials were then summed, and the resulting electric potential at a nerve cross section at the location of the stimulating electrodes was used to generate amplitude maps. For the amplitude modulation maps, the potential generated by each stimulation source was modulated with a sinusoidal signal with the respective carrier frequency, and the two potentials were then summed together to create a time-varying IS electric potential profile. The resulting signal at each spatial location was processed to extract the magnitude of amplitude modulation. The absolute value of the Hilbert transform was used to extract the envelope of the signal, and the envelope peak-to-peak amplitude was used to quantify the magnitude of amplitude modulation. To discretize the amount of amplitude modulation into three levels (high, intermediate, low), the amplitude modulation magnitudes at each fascicle location were extracted to generate the values distribution across the whole fascicle population. The three levels were then defined using the values quantiles as follows: low (values below quantile 0.33), intermediate (values between quantiles 0.33 and 0.66) and high (values above quantile 0.66).

#### Analysis of computational modeling results

The computational modeling outcomes include activation thresholds and voltage transient profiles for each fiber/stimulation parameter combination. This allows us to determine the presence of an action potential as a result of electrical stimulation. All analyses done on the fiber activation were performed using Python 3.9.

To investigate the correlation between physiological responses and fiber activation profiles obtained using the computational model, we performed a fascicle clustering based on the percentage of fiber types within each fascicle. We considered all fascicles that contained a majority of BP fibers to be BP-fascicles, while fascicles that had a prevalence of RL fibers were assigned to RL. Values of 6 μm and 10 μm were finally chosen to simulate the responses of slow and fast fibers respectively, as these values provided the best match with experimental data. BP fascicles were populated with a single, slow A-fiber (6 μm), while RL fascicles were populated with a single, fast A-fiber (10 μm). The knowledge of the function of fibers within the nerve acquired through fiber tracking (see Figure 2) allowed to simulate the outcome of electrical stimulation on target physiological functions. We considered the generation of an action potential (due to suprathreshold stimulation) for a fiber an indication of the effect on the respective physiological function. We computed the strength of the functional response as the percentage of relevant fibers being activated for specific physiological function, as described in (6).

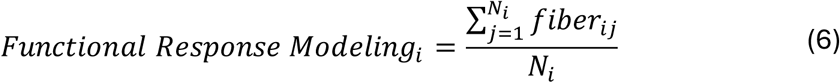

Here, *i* indicates the fiber population (BP or RL) with size *N*_i_, and *fiber_ij_* is a Boolean value indicating if an action potential was elicited (1) or not (0). The functional response (6) was computed for both sinusoidal and i^2^CS waveforms with a beat duration of 0.25 ms and for all steering ratios, similarly to in vivo experiments (see Figure 3). Functional responses obtained through modeling were directly compared with the respective physiological responses obtained experimentally for the same animal during current steering. A linear least-squares regression model was computed between the normalized (min-max normalization into the range of 0 to 1) functional responses obtained through modeling and the respective normalized functional responses obtained experimentally.

The functional responses derived from modeling were also used to compare the selectivity of sinusoidal stimulation and i^2^CS using the same approach deployed for experimental data. The functional responses as defined in (6) obtained at different steering ratios were fitted using a sigmoidal function as described in (3), and a selectivity factor was computed based on the parameters of the sigmoidal function as detailed in (4).

To study the fiber activation timing with respect to current steering and waveform type, we performed twofold analysis leveraging the ASCENT voltage transient protocol which allowed to obtain the timings of elicited action potentials. Firstly, we considered a single fast A-fiber (10 µm diameter) placed in a deep fascicle as displayed in Figure 6, inset i1. A single fiber type was selected to isolate the influence of current steering and waveform type on the timing of action potential. A total stimulation current of 2 mA was selected since it resulted in the generation of an action potential at all steering/waveform combinations. This allowed to study the impact of these stimulation parameters on the timing. Secondly, we considered all the RL-rich fascicles, which were populated with a single fast A-fiber (10 µm diameter), and we evaluated the activation timing resulting from different waveforms and steering ratios. This experiment was used to compare the laryngeal EMG responses obtained experimentally as a result of different waveforms and steering ratios with the responses obtained using the computational model for the RL-rich fascicles.

A similar approach was used to investigate the effect of steering/waveform selection on the spatiotemporal activation patterns for all the fascicles in the nerve model. We considered a single fast A-fiber placed in each of the nerve fascicles, during electrical stimulation with a total current of 2 mA. This resulted in most of the fibers being activated at each steering/waveform combination, allowing to investigate the effects of these stimulation parameters on the timing, as shown in Figure 6.

## Supplementary Material

**Figure S1:**
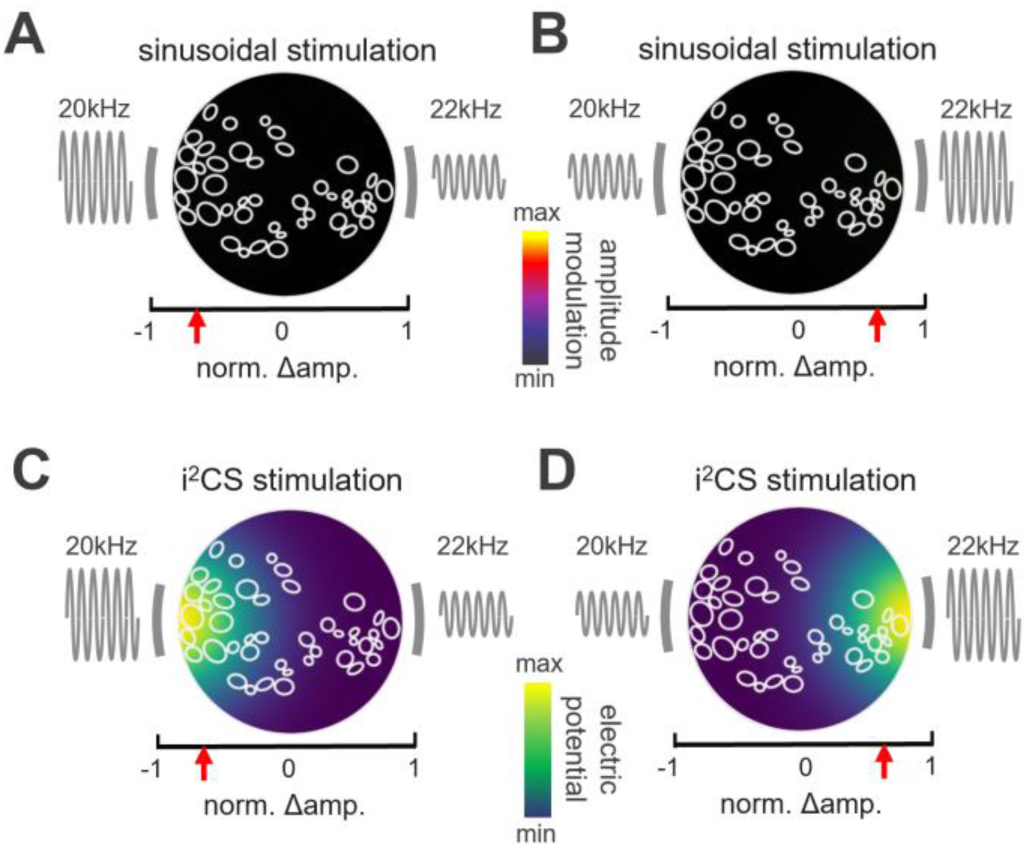
Interferential profile and electric potential obtained via sinusoidal stimulation and i^2^CS. (A) Sketch of the cross-section of the nerve, scaled fascicles from the real experiment are outlined in white. The colormap represents the peak-to-peak amplitude of the beat interference envelope as obtained during sinusoidal stimulation. (B) Same as in A, but for the other steering direction. (C) Same as in A), but the colormap represents the strength of the electric potential created by the 2 sources (grey bars) under a particular current steering ratio (red arrow on x-axis) during i^2^CS. The carrier frequencies and relative amplitudes are depicted next to the 2 sources. (D) Same as in C but for the other steering direction.

**Figure S2:**
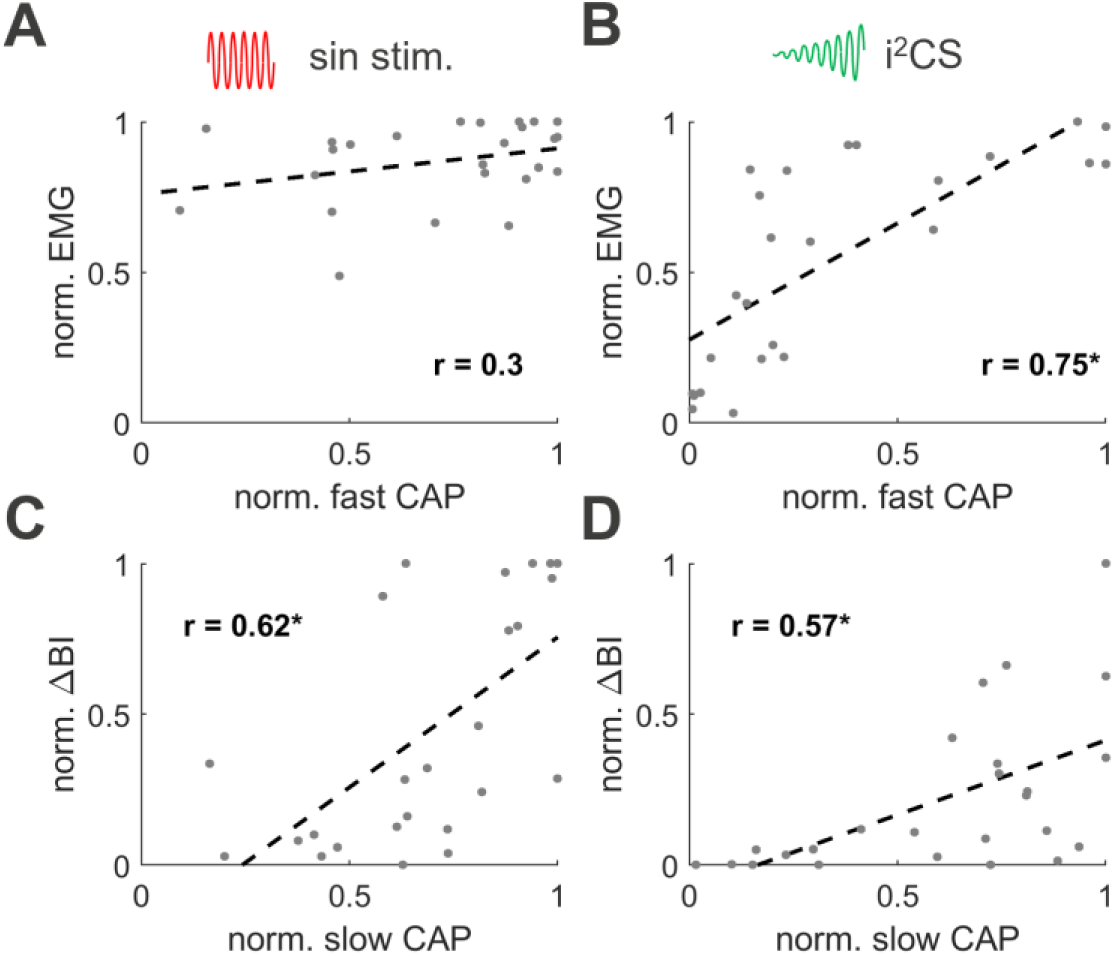
Correlation of eCAP and physiological responses. (A) Linear correlation between the fast A -fiber CAP response and the laryngeal EMG response upon sinusoidal stimulation (regression is denoted by dashed line, a star on the r-value denotes a significant correlation at p<0.01). (B) Same as in A, but for i^2^CS stimulation. (C) Same as in A, but for the linear correlation between the slow A-fiber CAP response and the breathing interval reduction. (D) Same as in C, but for i^2^CS stimulation. Datapoints in each panel represent responses of 5 different steering ratios from the same 5 animals depicted in Figure L (i.e., n = 25 in total).

**Figure S3:**
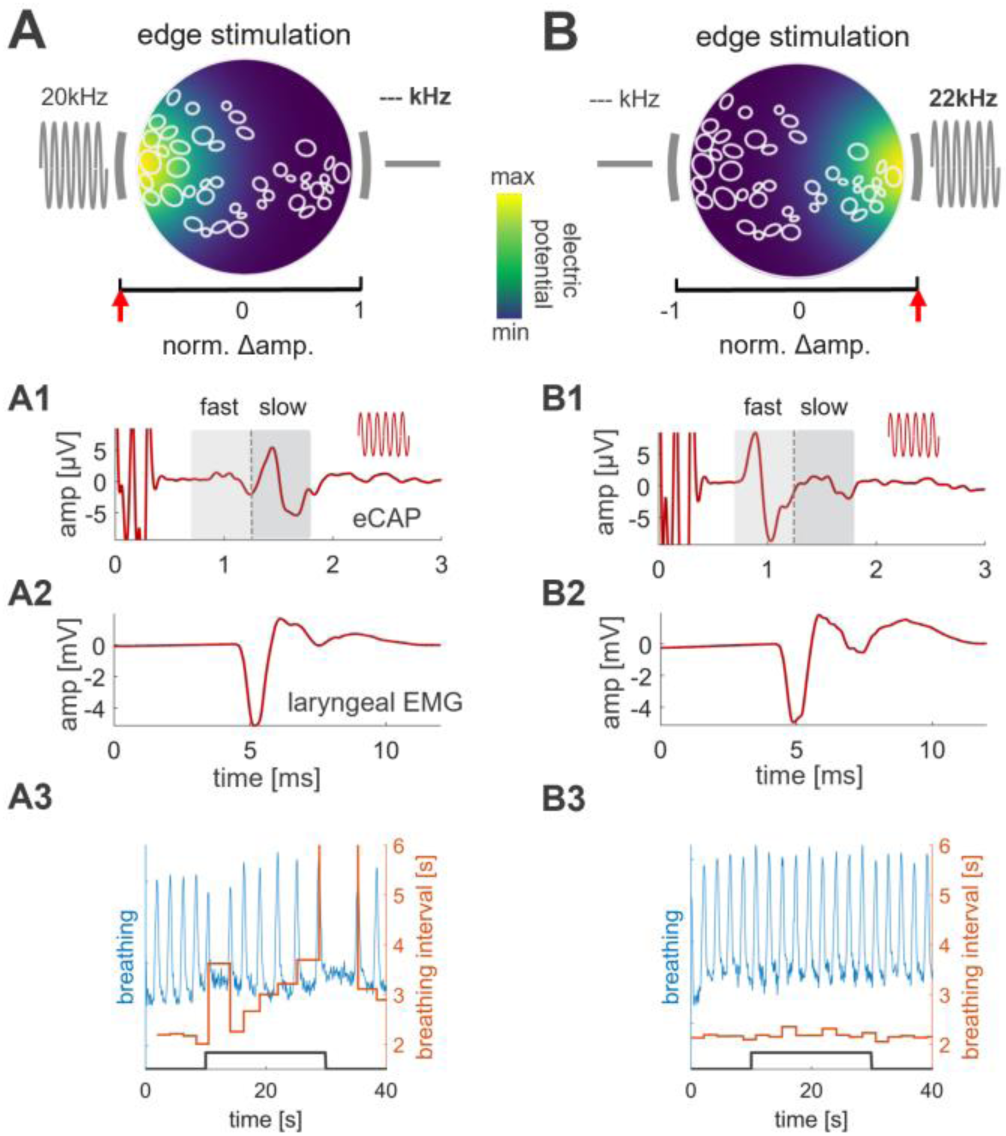
Pure sinusoidal edge stimulation. (A) Schematic cross section of a stimulated VN at the level of an implanted MCE; shown are outlines of nerve fascicles and the 2 contacts (grey bars) used for stimulation, with the left source at maximum amplitude (red arrow on left side of x - axis) and the right source inactive; The colormap represents the strength of the electric field. (A1) eCAP response, triggered from the onset of stimulation, with 1.5 mA total current delivered through the left source only; slow and fast eCAP components are identified by the shaded areas corresponding to time windows defined by the average conduction velocities for ‘slow’ and ‘fast’ A -fibers. (A2) Strong laryngeal EMG response to edge stimulation. (A3) Breathing response (blue) and respective change in breathing interval (orange during a 20 sec long train of edge stimulation (black trace). (B) Same as in A, but for the opposite electrode contact (i.e., the right side). The laryngeal EMG response is similar to that for left side stimulation, whereas the breathing response is absent.

**Figure S4:**
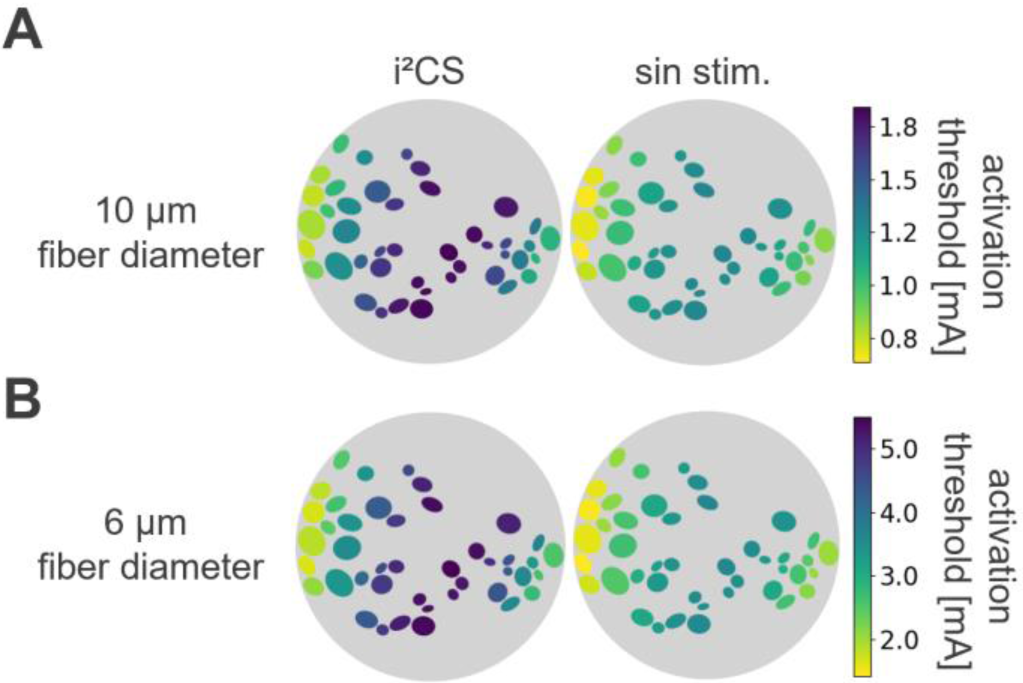
Activation thresholds for different fiber diameters and stimulation conditions. (A) Activation threshold for 10 μm fibers for i^2^CS (left) and sinusoidal stimulation (right) at a steering ratio of 0, showing an increased threshold at the center of maximum AM for i^2^CS compared to sinusoidal stimulation. (B) Same as in A but for 6 μm fibers. Note the similarity in relative activation thresholds for different fascicles between A and B.

**Figure S5:**
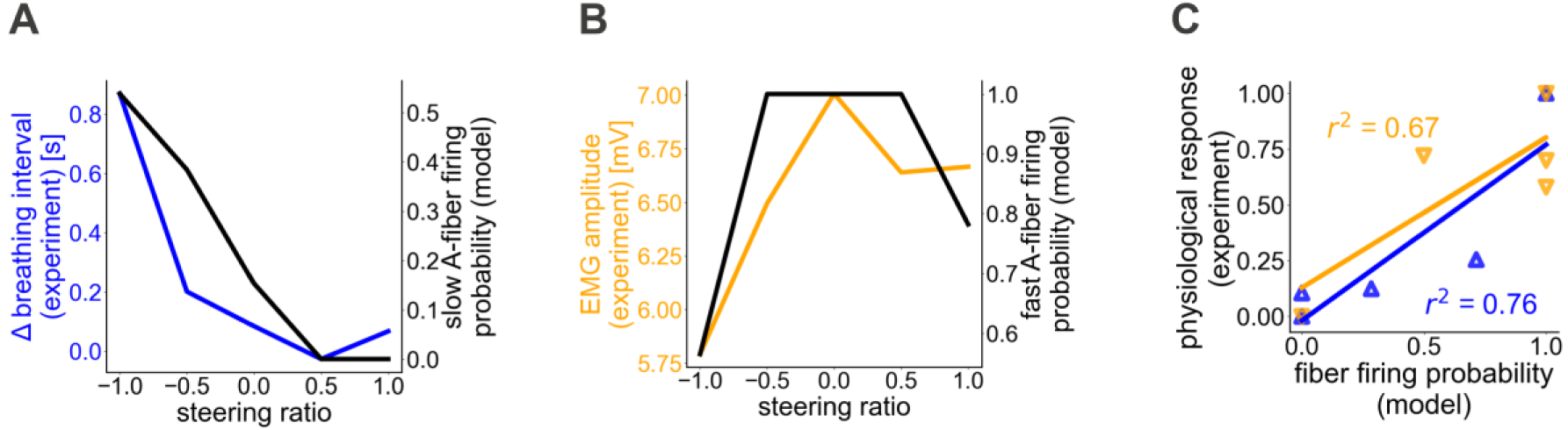
Anatomically realistic biophysical model of the nerve -electrode interface replicates experimentally measured activation of organ-specific fibers in response to sinusoidal stimulation. (A) Modeled slow A-fiber responses to sinusoidal stimulation with different steering ratios (having combined total current amplitude of 1.5 mA) and change in breathing rate experimentally measured using sinusoidal stimulation with the same steering ratios. (B) Modeled fast A-fiber responses and laryngeal EMG responses recorded experimentally. (C) Correlation between modeled fiber firing probabilities and normalized physiological responses obtained experimentally in the same animal: fast A-fibers vs. laryngeal EMG (orange), slow A-fiber vs. breathing response (blue).

**Figure S6:**
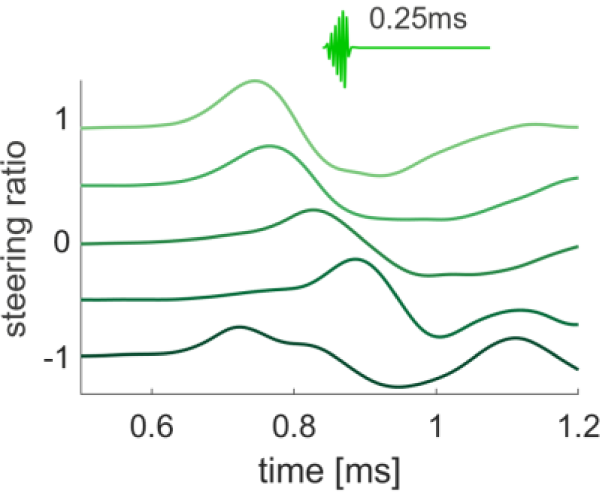
CAP responses from one example animal to a 0.25 ms long i^2^CS stimulation at different steering ratios (total amplitude 2 mA).

**Figure S7:**
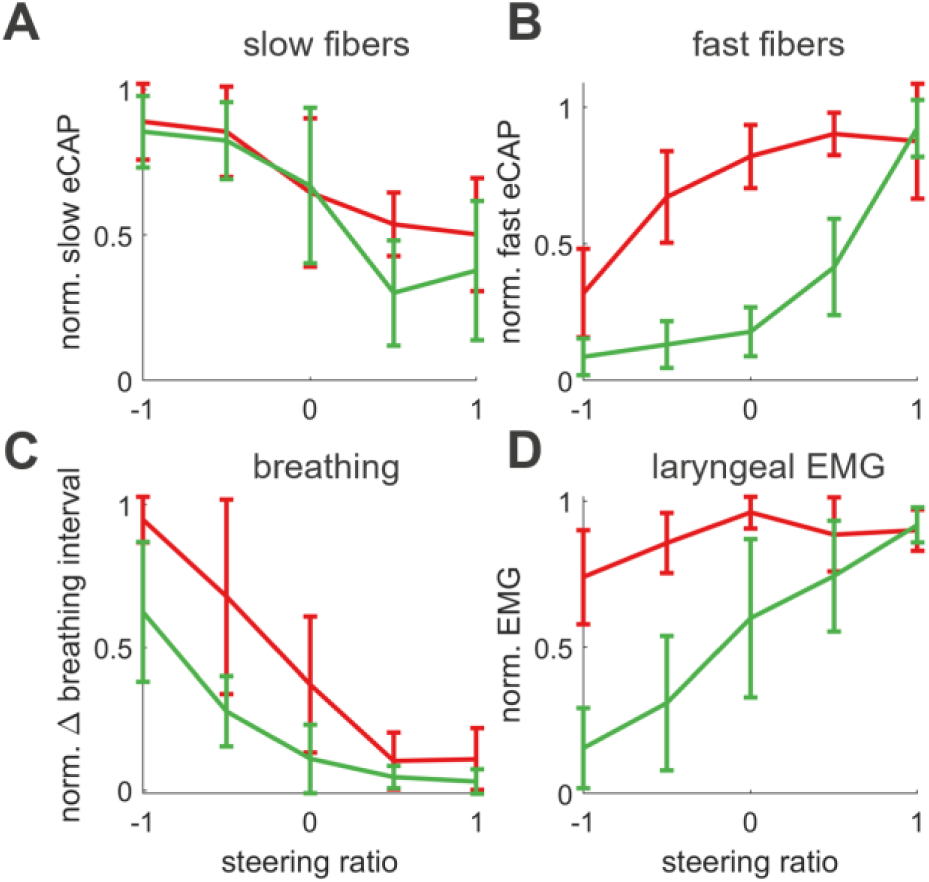
Normalized eCAP and physiological response across the population indicates a differential effect of steering ratio between i ^2^CS and sinusoidal stimulation. (A) Normalized slow eCAP responses for sinusoidal and interferential stimulation at different steering ratios. (B) Same as in (A), but for fast eCAPs. (C) Normalized magnitude of the (desired) breathing response (change in breathing interval) at different steering ratios, for interferential and equivalent sinusoidal stimulation. (D) Normalized amplitude of the (undesired) laryngeal EMG at different steering ratios. All data is shown as mean±SD from n = 5 animals.

**Figure S8:**
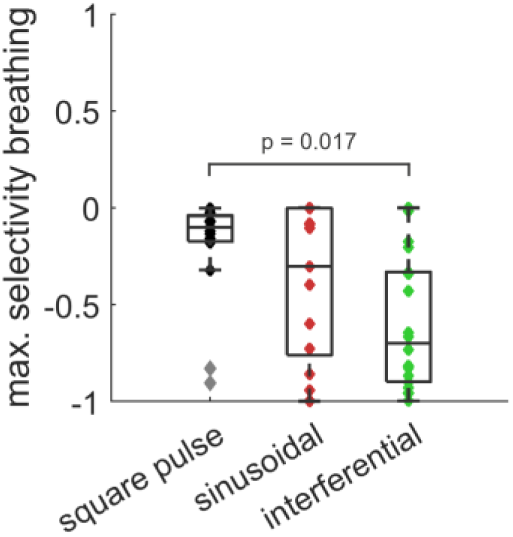
Comparison of square pulse, sinusoidal and interferential stimulation selectivity. Selectivity indices (SIs) for pulsed stimulation extracted from functional mapping data reported in (Jayaprakash et al., 2023). The electrode with the largest selectivity value for each stimulation strength was used; only stimulus trains eliciting >20% BI reduction from at least 1 electrode were considered, resulting in 20 observations in 4 animals. Sinusoidal and interferential data were extracted from newly performed experiments (6 animals, 30 datapoints).

**Figure S9:**
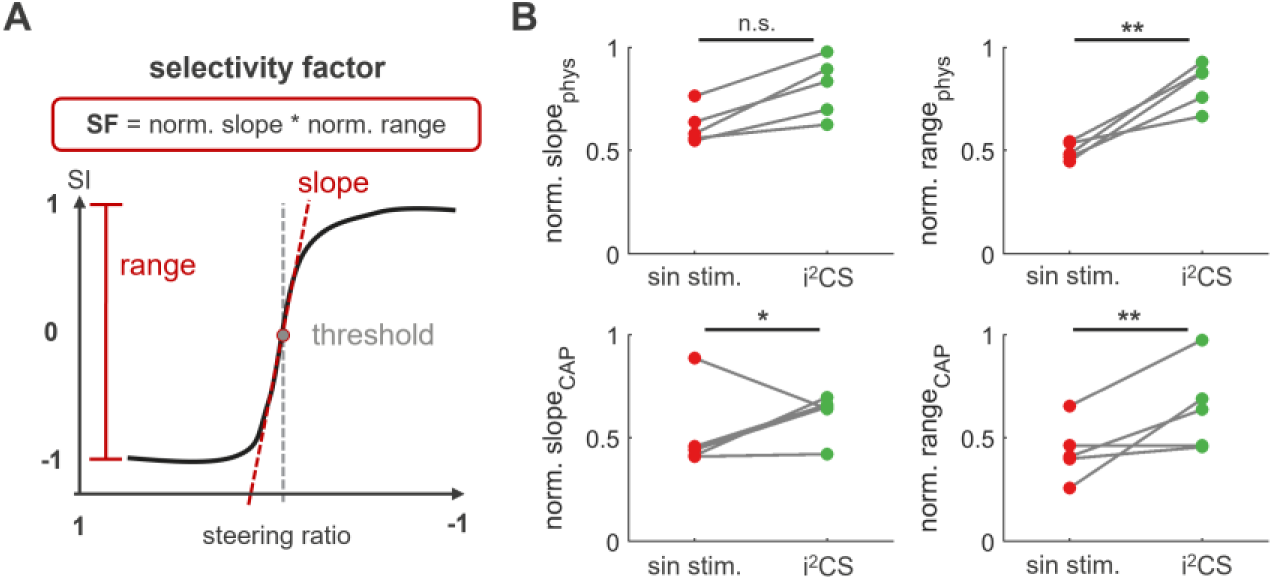
Calculation of selectivity factor and underlying parameters. (A) The selectivity factor is calculated from two key parameters of the sigmoidal fit to the data: The slope can be seen as a measure of how sharp the separation between the two readouts is in terms of spatial selectivity and is calculated as the normalized angle between the slope and the x-axis. The range represents a measure of the maximal, relative difference between the two readouts in terms of selectivity and is calculated as the normalized difference between the maximum and the minimum value of the function. Finally, the threshold is the x-axis value where the slope is maximal and provides spatial information about where on the cross -section the shift in selectivity occurs. (B) All values from A plotted separately for all animals used in the comparison. Note that most parameters show significantly higher values for interferential stimulation (*p<0.05 **p<0.01, Wilcoxon rank-sum test).

**Figure S10:**
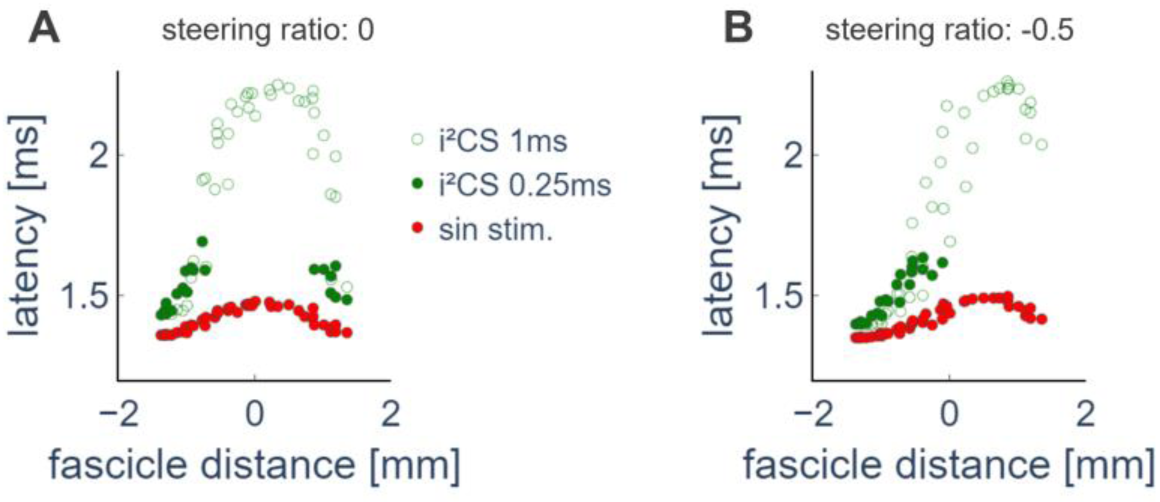
Spatio-temporal activation pattern within the nerve during sinusoidal stimulation and i^2^CS with different beat durations. (A) Time of fiber activation for each A -fiber based on the respective fascicle index resulting from the application of different stimulation waveforms for a current steering in the center of the nerve (steering index = 0) and a total amplitude of 1.5 mA: PS0 (red), i^2^CS with a beat duration of 0.25 ms (green dots), and i^2^CS with a beat duration of 1 ms (green empty dots) (B) Same as A), but for a current steering towards the left side of the nerve cross-section (steering index = −0.5).

**Figure S11:**
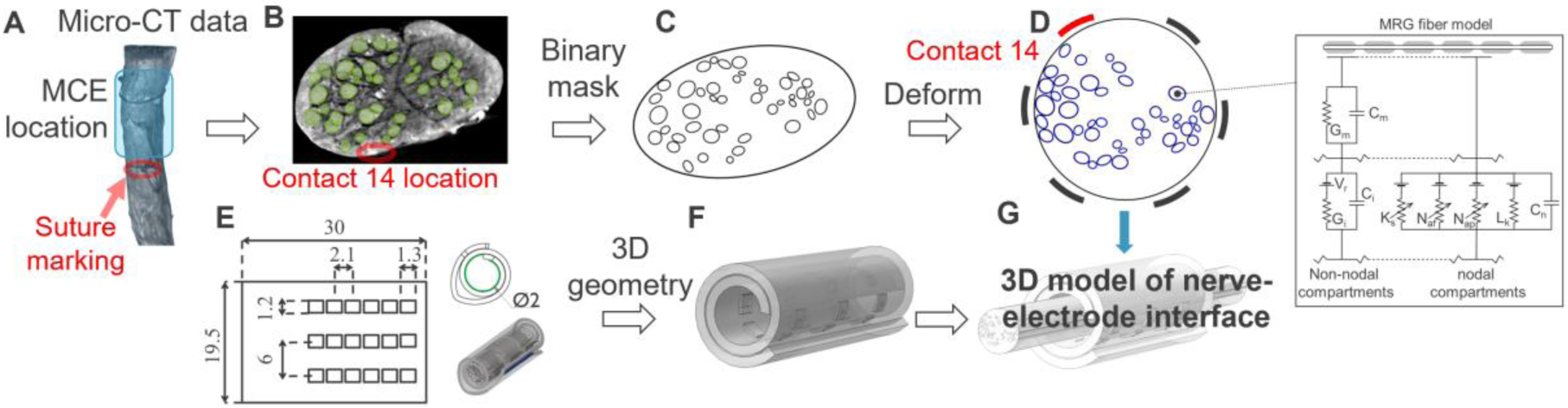
Pipeline for computational modeling. (A-D) Histological images obtained from the VN at the cuff location are analyzed to extract the boundaries of the whole nerve and single fascicles. The binary mask containing these boundaries is then deformed to a circular shape to account for the deformation resulting from cuffing the nerve. (E, F) The geometrical parameters of the cuff are used to derive a 3D model of the cuff. (G) The 3D nerve model and cuff geometry are combined to obtain the final 3D model.

**Figure S12:**
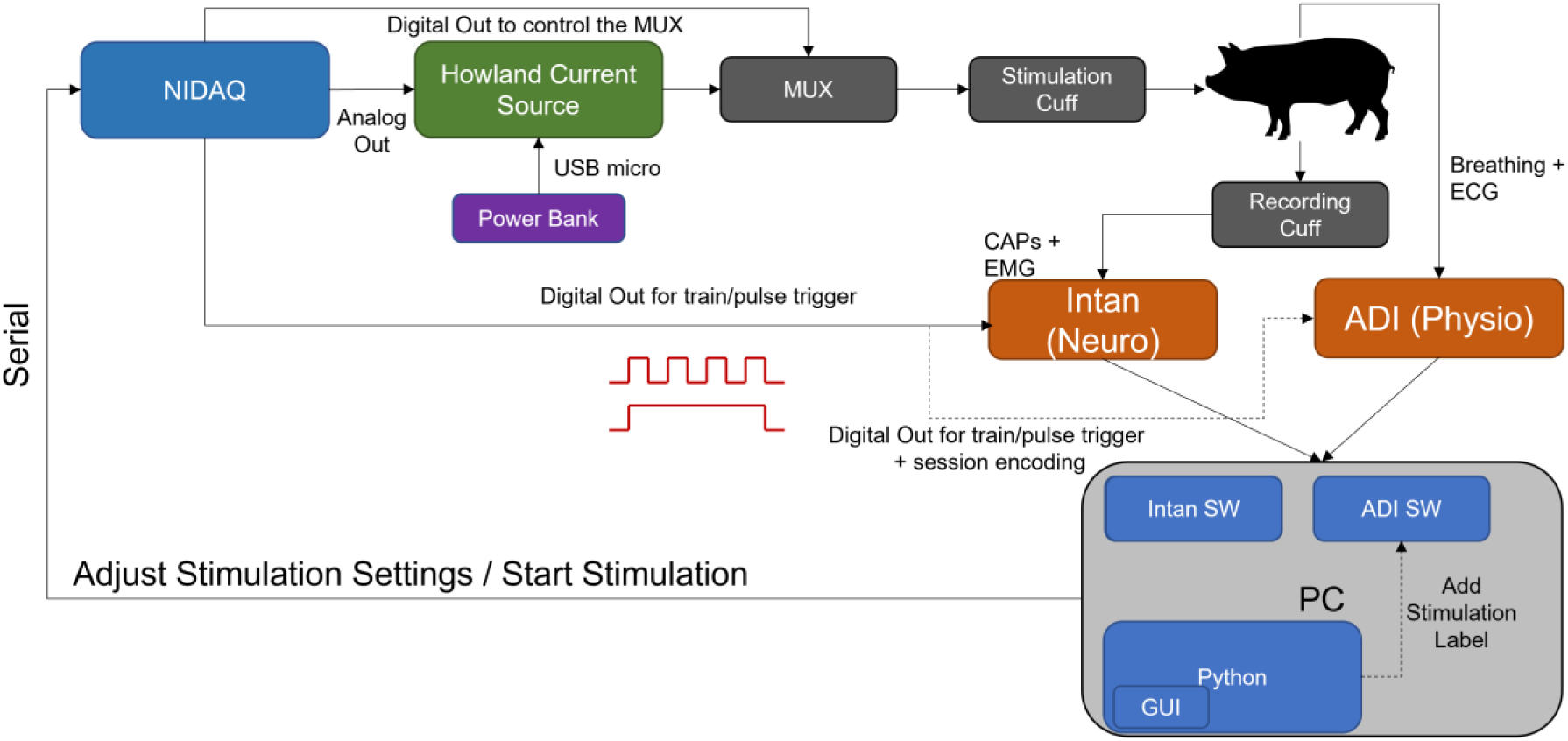
Diagram of the experimental setup for in-vivo experiments. The PCIe-6363 DAQ is controlled via a PC to produce the required analog stimulation waveforms and generates additional digital signals used to mark the start and end of each pulse and of the whole stimulation train. The voltage signals are transmitted to the custom Howland current source to convert the signal to current. The current signals are then multiplexed to address specific electrodes on the stimulation cuff. Neural and physiological signals are recorded from the animal and transmitted to the PC.

**Figure S13:**
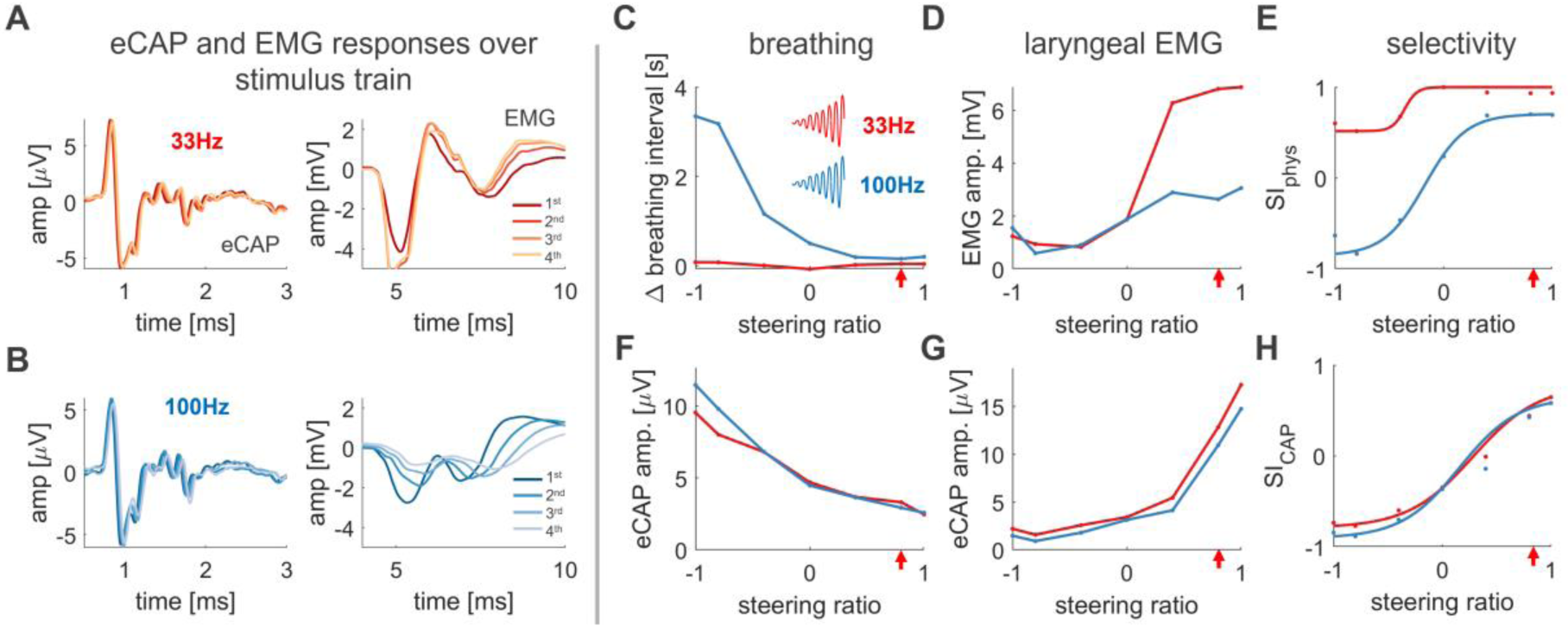
Effect of repetition frequency on breathing and laryngeal EMG response. (A) Example average eCAP (left) and EMG (right) responses for the first (darkest trace) to last (lightest trace) quarter of the 20 sec stimulus train to 33 Hz inter -stimulus-interval (ITI) i^2^CS stimulation at a steering ratio of 0.9. (B) Same as in A, but for 100 Hz ITI. Note the reduced laryngeal EMG response over time while the eCAP response is unaffected. (C) Breathing interval at different steering ratios. Note the much stronger activation of breathing responses at 100 Hz ITI. (D) Average amplitude over the whole stimulus train of the laryngeal EMG response at different steering ratios. (E) Selectivity index as calculated from the physiological responses in C and D, fitted with a sigmoidal function. (F) eCAP amplitude for slow A-fibers at different steering ratios. (G) Same as in F, but for fast A-fibers. (H) Selectivity index as calculated from eCAP amplitudes in F and G, fitted with a sigmoidal function. Red arrows denote the steering ratio of the example responses shown in A and B.

